# *In situ* structure determination of Respiratory Supercomplexes and ATP synthase oligomers in mammalian mitochondrial inner membrane

**DOI:** 10.1101/2025.09.19.677273

**Authors:** Atsuki Nakano, Takahiro Masuya, Shinsuke Akisada, Moe Ishikawa-Fukuda, Kaoru Mitsuoka, Hideto Miyoshi, Masatoshi Murai, Ken Yokoyama

## Abstract

To understand how membrane protein complexes function within biological membranes, it is essential to determine their structure in their natural membrane environment. Here, we employed cryoEM structure analysis to elucidate the structures of ATP synthase F_o_F_1_ and respiratory Supercomplexes (SCs) on sub-mitochondrial particles (SMPs) isolated from bovine heart mitochondria. On SMPs, the majority of F_o_F_1_ was identified as dimers bound by the regulatory factor dimeric IF_1_. In addition, a tetrameric structure formed by association of F_o_F_1_ IF_1_ dimers and with a linear arrangement of the F_1_ head were also identified. These structures induced a steep membrane curvature, indicating the presence of a structure on SMPs similar to that found on the tips of mitochondrial cristae. High-resolution structures of the respiratory complexes were also determined, and sub-class structures of both CI and CIII_2_ were resolved. Most SCs were of the CI_1_CIII_2_CIV_3_ structure, although the presence of the CI_2_CIII_2_CIV_6_ mega complex was also identified. Our study enabled rapid *in situ* structural determination of SCs and F_o_F_1_ ATP synthase from small amount of membrane fractions, paving the way for elucidation of the molecular basis of metabolic disorders and mitochondrial diseases at the level of higher-order architecture.

## Introduction

Membrane proteins play essential roles in cellular functions, including signal transduction, energy conversion, and molecular transport across membranes ^1,2^. Structural analysis of membrane proteins typically involves detergent-based extraction from biological membranes, followed by multiple purification processes. These structures of membrane protein are then determined using X-ray crystallography or cryo-electron microscopy (cryo-EM). However, during solubilization and purification, loosely bound subunits may dissociate, and essential lipids might be replaced by detergents. Additionally, higher-order super molecular structures formed by the association of multiple membrane protein complexes are likely to be lost during purification processes. Consequently, the structure of solubilized membrane proteins is likely to differ from *in situ* structure of membrane proteins in their native biological context.

Mitochondria, intracellular energy factories, are complex organelles composed of an outer membrane and an inner membrane. The inner membrane hosts the respiratory complexes and ATP synthase, which sustain life by synthesizing ATP through the multistep process of oxidative phosphorylation^3^.

The mammalian mitochondrial respiratory system comprises four enzyme complexes, designated complexes I, II, III, and IV. Complex I (NADH-ubiquinone oxidoreductase, CI) functions as a primary entry point for electrons from matrix-derived NADH, catalyzing the reduction of ubiquinone (UQ) to ubiquinol (UQH_2_)^4-7^. Complex II (succinate-ubiquinone oxidoreductase, CII) provides an additional entry point for electrons through the oxidation of succinate to fumarate in the tricarboxylic acid (TCA) cycle, using UQ as the electron acceptor^8,9^. UQH_2_ then donates electrons to complex III (ubiquinol-cytochrome *c* oxidoreductase, CIII), which reduces cytochrome *c* (cyt. *c*) on the intermembrane space side of the inner membrane^8,9^. Electrons are subsequently passed to complex IV (cytochrome *c* oxidase, CIV), where molecular oxygen—the terminal electron acceptor—is reduced to water^10,11^. Electron transfer through complexes I, III, and IV is coupled to proton translocation from the matrix to the intermembrane space, generating an electrochemical proton gradient—known as the proton motive force (*pmf*)—across the inner membrane, which drives ATP synthesis via the F_o_F_1_-ATP synthase.

These respiratory chain complexes (I-IV) were first solubilized and purified separately from the mitochondrial inner membrane, and their crystal or cryoEM structures have been determined^12-15^. Recently, gentle solubilization of the inner mitochondrial membrane revealed that complexes I, III, and IV associate to form larger Supercomplexes (SCs), whose cryoEM structure has also been determined^16^.

ATP synthase F_o_F_1_ utilizes the *pmf* generated by the respiratory chain complexes to rotate the hydrophobic rotor *c*_*8*_-ring of the F_o_ domain relative to the stator complex^3,17,18^. This rotation drives the phosphorylation of ADP to synthesize ATP in the F_1_ domain. The bovine mitochondria F_o_F_1_, initially purified as a monomer^19^, has been purified as a dimer through careful solubilization and purification, and its cryo-EM structure also determined^20^. Single-particle analysis of porcine mitochondrial inner membrane fractions solubilized with detergents has reported a tetrameric structure of F_o_F_1_, suggesting that multiple F_o_F_1_ molecules associate and function as a super molecular complex in the mitochondrial inner membrane^21^.

Mutations within genes encoding the respiratory complex and F_o_F_1_ ATP synthase subunits are frequently associated with mitochondrial disorders and metabolic diseases such as diabetes, highlighting their significance as potential therapeutic targets^22,23^. Yet, the link between disease pathology and the higher-order architecture of these complexes remains poorly understood. Cryo-electron tomography (cryo-ET) offers a powerful approach for visualizing macromolecular structures within the context of organelle and cellular membranes. Nevertheless, the time-intensive acquisition of tilt series and the complexity of three-dimensional reconstruction have thus far limited its ability to resolve fine substructures of membrane-embedded proteins at high resolution.

In this study, we determined structures on the inner mitochondria membranes through direct observation of sub-mitochondrial particles (SMPs) using cryo-EM single particle analysis. For the respiratory SCs we obtained structures with resolution comparable to that of the individual solubilized and purified complexes. We also elucidated the structure of the dimeric and tetrameric F_o_F_1_ and obtained a high-resolution structure of the monomeric form, which can be used to construct an atomic model.

## Results

### Preparation of bovine heart SMPs and data acquisitions respiratory SCs and F_o_F_1_ oligomer

Submitochondrial particles (SMPs) were prepared from bovine heart mitochondria^24^ (Figure 1A). The SMPs, suspended at a concentration of 7 mg protein/ml in a buffer containing 50 mM sucrose and 10 mM Tris/HCl (pH 7.4), were blotted onto a quantifoil UltraAufoil grid and flash-frozen in liquid ethane. Cryo-EM micrographs were acquired with a Titan Krios equipped with a K3 detector. A typical cryo-EM image is shown in Figure 1B. The F_1_ domain of F_o_F_1_ can be clearly observed in the micrograph where the ice thickness is relatively thin (Figure 1B, lower panel). First, we reconstructed the 3D structures of both the F_o_F_1_ dimer and SCs from 5k micrographs, which serves as machine learning training data for subsequent particle picking. From approximately 30,680 movies, ∼4 x 10^6^ single-particle images of F_o_F_1_ dimer and SCs were extracted and subjected to further single-particle analysis (Figure S1A and B).

**Figure 1.**
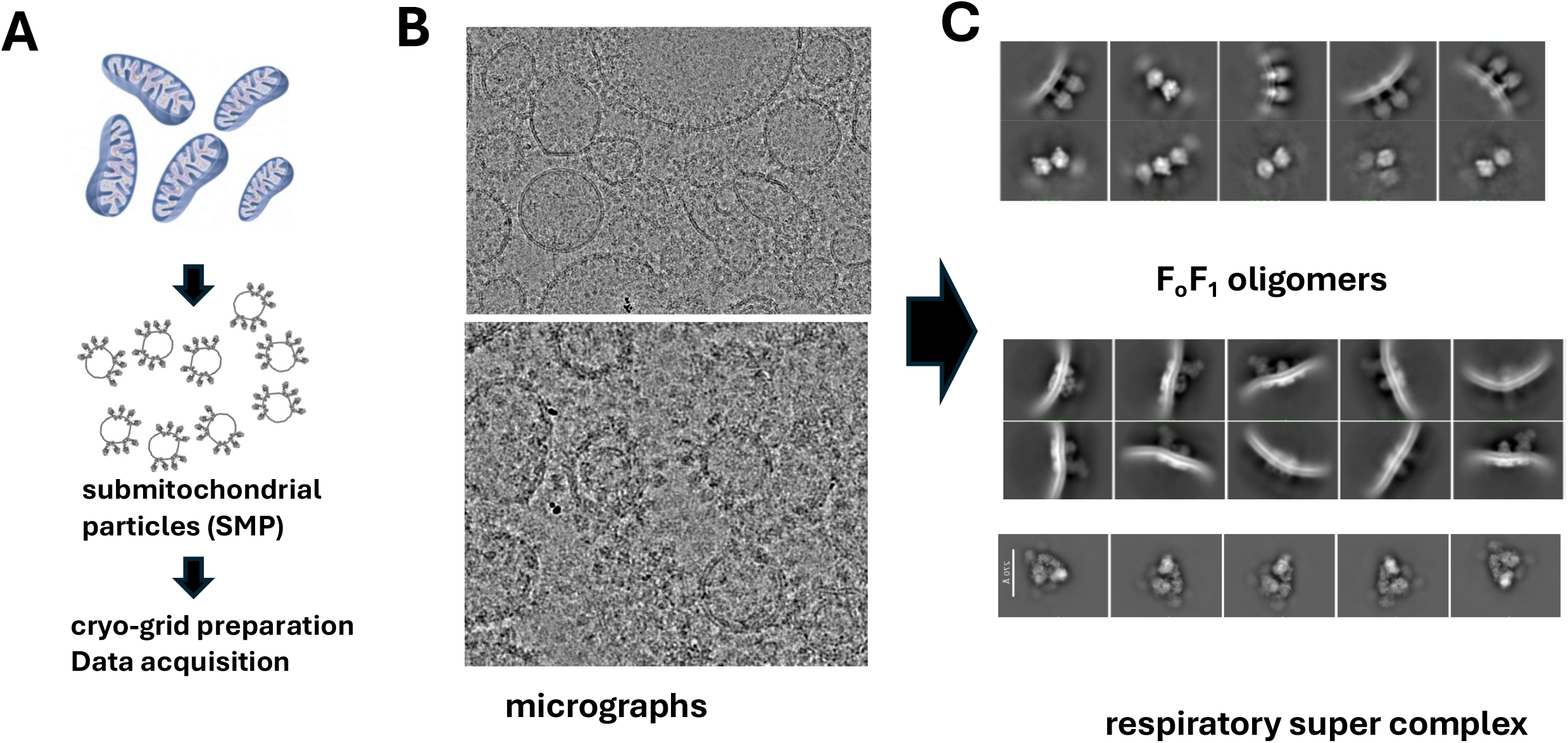
Cryo-EM imaging of SMPs and representative 2D class averages of F_o_F_1_ oligomers and Supercomplexes (SCs). **(A)** Schematic illustration of SMP preparation from disrupted bovine heart mitochondria. Details of SMP preparation are described in the Materials and Methods section. **(B)** Cryo-EM micrograph of isolated SMPs (bottom, magnified view). **(C)** Representative 2D class averages of F_o_F_1_ oligomers (top) and SCs on SMPs viewed from the side (upper) and top (lower).

### 3D reconstruction of the F_o_F_1_ oligomer in the SMPs

Using two adjacent F_o_F_1_ dimers as templates, Topaz, a machine learning-based tool, selected 1,306 k particles from ∼4 x 10^7^ particles including both F_o_F_1_ dimer and SCs (Figure S1A). After homogeneous refinement, a structure of F_o_F_1_ dimers connected laterally by a rod-shaped molecule, likely IF_1_, was obtained from 382 k particles (Figure S1B and 2A). Hereafter, the F_o_F_1_ dimer connected by IF_1_ is referred to as the F_o_F_1_–IF_1_ dimer, with its left and right protomers designated F_o_F_1_–1 and F_o_F_1_–2, respectively.

The resolution of the F_o_F_1_-IF_1_ dimer structure is limited by the fluctuations of the two monomers relative to one another, with an FSC resolution of approximately 7Å. The F_o_ domain of the F_o_F_1_-IF_1_ dimer is embedded within the lipid bilayer (Figure 2A). On the matrix side, density corresponding to a domain of the *c*_*8*_-ring and the *e*-subunit (indicated by red and blue arrows, respectively), which is connected to the *c*_*8*_-ring, was clearly observed.

**Figure 2.**
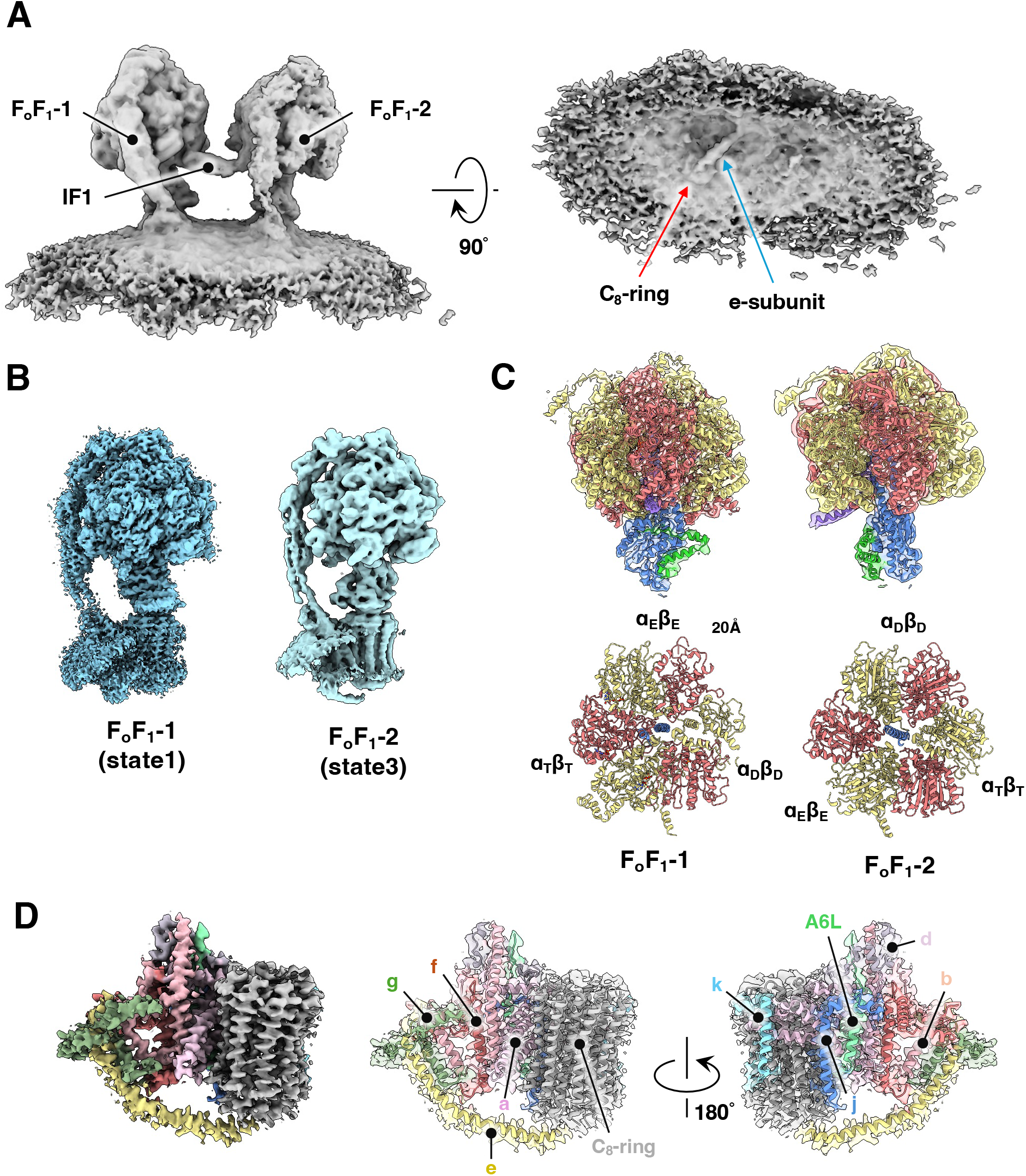
Cryo-EM structures of the F_o_F_1_-IF1 dimer in the SMP membrane. (**A**) Cryo-EM map of the F_o_F_1_–IF_1_ dimer, shown in side view (left panel) and from the matrix side of the membrane (right panel). In the matrix side view, the *c*_*8*_-ring and the *e* subunit are clearly visible. (**B**) Focused refinement maps of the two F_o_F_1_ monomers that constitute the F_o_F_1_–IF_1_ dimer: F_o_F_1_–1 and F_o_F_1_–2 correspond to rotational state 1 and rotational state 3, respectively. (**C**) Atomic models of the F_1_ domain from F_o_F_1_–1 and F_o_F_1_–2. Side views (top panel) and top views (lower panel) are shown. In F_o_F_1_–1, nucleotides were modeled: no density was observed at the α_E_β_E_ catalytic site, whereas ATP–Mg^2+^ and ADP were bound at α_T_β_T_ and α_D_β_D_, respectively. In F_o_F_1_–2, nucleotide modeling was not performed due to resolution limitations. (**D**) The focused refinement map of the F_o_ domain from F_o_F_1_–1 (left panel), and the corresponding atomic model (right panel), with each subunit shown in a distinct color.

To obtain a high resolution structure the monomeric F_o_F_1_, masked structural refinements were carried out on each monomer, (Figure S1B). The F_o_F_1_-1 and F_o_F_1_-2 structures were determined at resolutions of 3.3 Å and 3.7 Å, respectively (Figure 2B, S1C), allowing the construction of atomic models (Figure 2C).

The structure of the F_o_F_1_-IF_1_ dimer on the SMPs is very similar to the previously published structure of the F_o_F_1_-IF_1_ dimer unit which forms part of the solubilized tetrameric F_o_F_1_ from porcine heart mitochondria^21^. In each monomer, the F_1_ domain exhibited a distinct rotational state relative to the central axis. F_o_F_1_-1 was in rotational state 1, while F_o_F_1_-2 assumed rotational state 3 (Figure 2C). Using focused refinement on the membrane domain of F_o_F_1_, we obtained a map of the F_o_ domain of F_o_F_1_-1 at a resolution of 4.1 Å. In this structure, we identified not only the transmembrane helices of both the *c*_8_-ring and stator *a*-subunit but also the long helix of the *e*-subunit (Figure 2D, indicated by blue arrow), which connects the stator domain of F_o_ with the lipid in the central pore of the *c*_8_-ring (Figure 2D, indicted by red arrow).

Single-particle cryo-EM analysis of the solubilized fraction of porcine heart mitochondrial SMP demonstrated the presence of a tetrameric F_o_F_1_ structure^21^. To investigate whether a similar tetrameric structure exists in bovine heart mitochondrial SMPs, we extracted F_o_F_1_-IF_1_ dimer images with extended pixel size from micrographs (981 x 981 Å^2^) and reconstructed 3D images with an extended box size (Figure S1B). The resulting structures contained two sets of F_o_F_1_-IF_1_ dimers (Figure 3A). When viewed laterally, the oligomeric F_o_F_1_ exhibited a V-shaped membrane curvature, with the F_o_F_1_-IF_1_ dimers positioned at both sides of the apex of this roughly triangular curvature (Figure 3A, *lower*). This arrangement of the F_o_F_1_-IF_1_ dimer closely resembles the tetrameric F_o_F_1_ structure derived from porcine heart mitochondria. In the present analysis, densities corresponding to putative F_1_ structures were observed adjacent to the tetramer unit (Figure 3A, *right*). This finding suggests that, within SMPs, F_o_F_1_-IF_1_ dimers may assemble into a linear arrangement, potentially contributing to a crista-like structure.

**Figure 3.**
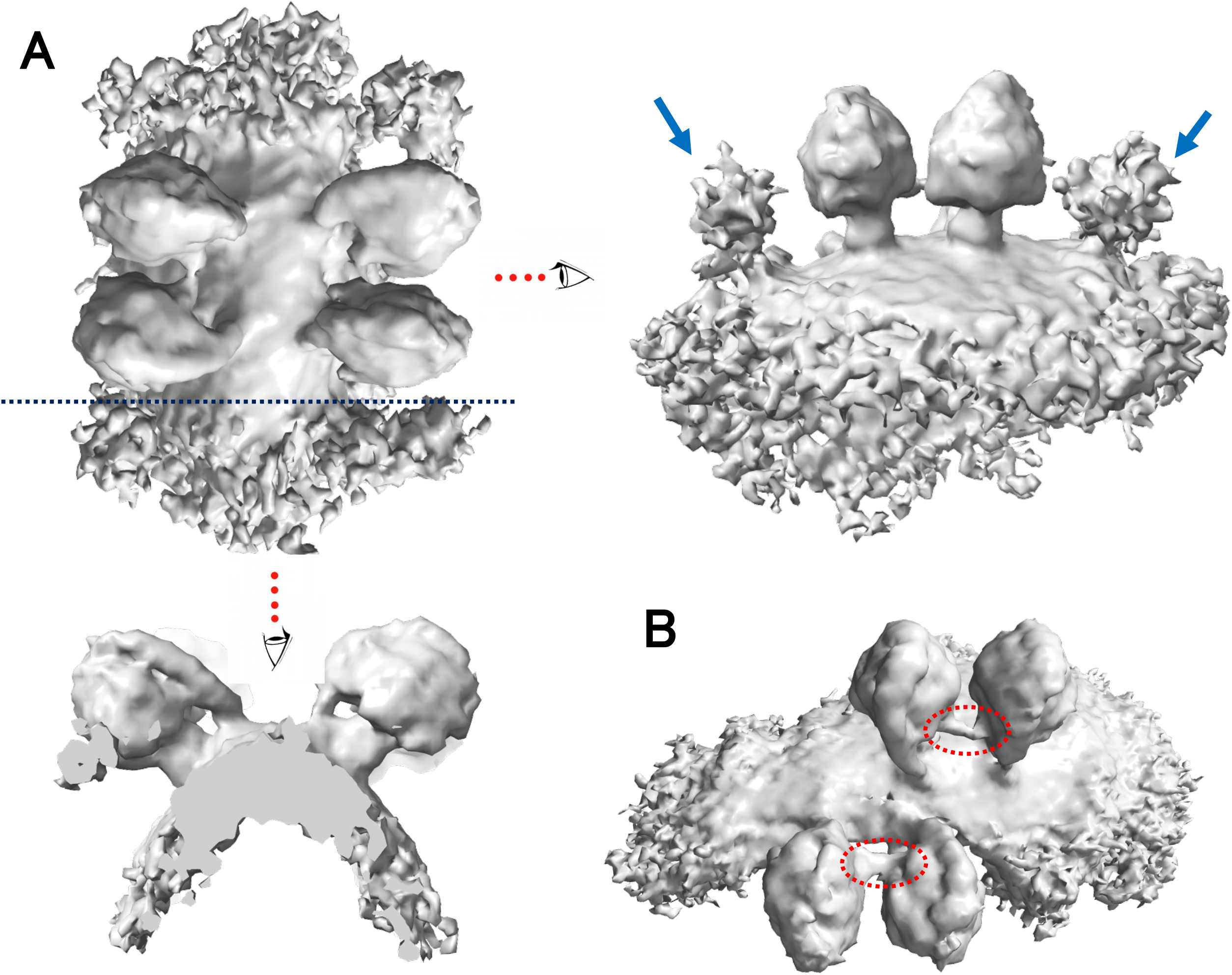
Cryo-EM structures of F_o_F_1_ tetramers on the SMP membrane. (A) Cryo-EM density map of the F_o_F_1_ tetramer. The upper panel shows the overall structure, while the lower panel presents a cross-sectional side view, and the right panel displays an intact side view. Putative F_1_-head domains are indicated by blue arrows. (B) Cryo-EM map of a refined subclass of the F_o_F_1_ tetramer obtained through 3D classification. Densities attributed to IF1 are highlighted with red ellipses.

Additionally, we identified another oligomeric F_o_F_1_ structure distinct from the tetrameric F_o_F_1_. In this oligomer, the arrangement of the F_o_F_1_-IF_1_ dimer across the apex of the curved membrane was shifted by one F_1_ unit (Figure 3B). Consequently, the state1 F_1_ domains of the IF1 dimers faced each other across the apex of the membrane. This result suggests that the binding between F_o_F_1_-IF_1_ dimers across the membrane apex is weak.

### 3D reconstruction of the Respiratory Supercomplexes in the SMPs

As described above, we performed four rounds of heterogeneous refinement on approximately 3 million particles containing a mixture of F_o_F_1_ oligomers and SCs, which resulted in the isolation of a subclass containing predominantly SCs, comprising 764,000 particles (Figure S1A). Homogeneous refinement of this subclass yielded a 3.5 Å-resolution structure of the SCs.

Subsequent classification allowed us to distinguish two distinct populations: one consisting of SCs composed of CI and the CIII dimer (CIII_2_), and another exhibiting an additional density adjacent to the distal end of CI (Figure S1A). As described later, this contact site corresponds to the ND5 subunit. This location of extra density coincides with that of CIV observed in previously reported SC structures from solubilized mitochondria^25^, suggesting that it represents CIV unit.

We then selected the class showing clear CIV density through heterogeneous refinement, followed by focused refinement on the CIV region using the resulting 227,000 particles. This processing yielded a 2.61 Å-resolution structure of CIV within the SC. In the SC density map, two additional CIV-like densities were observed adjacent to the CIII_2_, although they were less well-defined compared to the canonical CIV density (Figure 4A, inset). One was located in the interspace between CI and CIII_2_ (location B in the inset of Figure 4A), and the other was associated with solely CIII_2_ (location C in the inset of Figure 4A).

**Figure 4.**
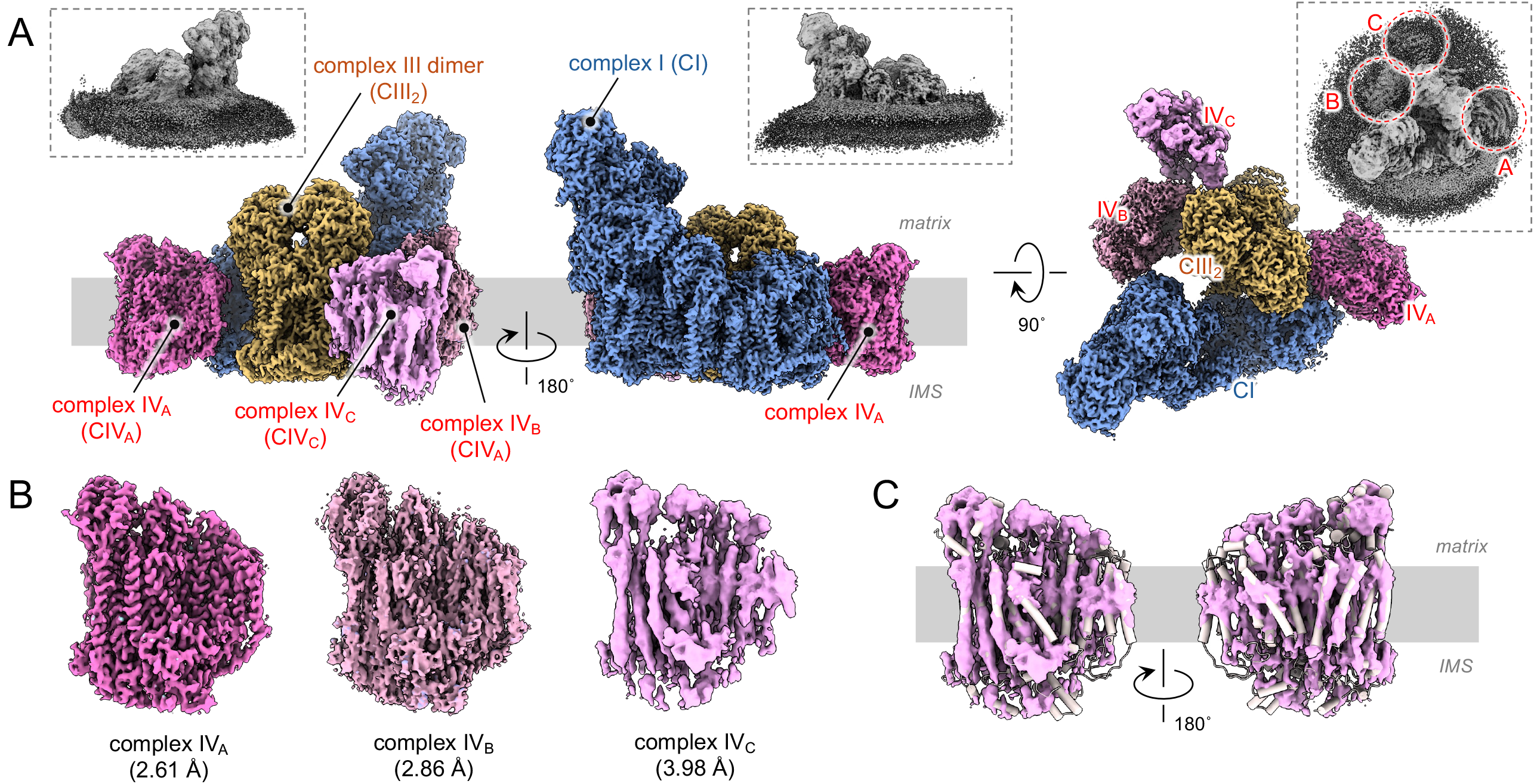
Structure and organization of the respiratory supercomplex (SC). (A) Overall architecture of the mitochondrial respiratory SC. Cryo-EM densities of five respiratory complexes—CI-1 (blue), CIII_2_-1 (yellow), CIV_A_ (deep pink), CIV_B_ (pink), and CIV_C_ (light pink)—are shown. The final composite map was generated by merging the individually refined densities, as outlined in Figure S1B. Insets display the EM density maps of SC after homogeneous refinement of 764,763 SC particles (3.5 Å, Figure S1B). (B)Comparison of EM maps corresponding to complexes CIV_A_, CIV_B_, and CIV_C_. (C) Superposition of the atomic model of complex IV, built from the CIV_A_ density, onto the CIV_C_ density map. This fitting confirms that the CIV_C_ density corresponds to complex IV. The model is shown in a tube helix representation to highlight its correspondence with the CIV_C_ map.

Local refinement of each CIV unit, combined with subtraction of surrounding densities and particle selection via 3D classification, resulted in improved maps featuring clearly resolved transmembrane helices (Figure 4B). The CIV associated with distal end of CI was designated as CIV_A_, the one located between CI and CIII_2_ as CIV_B_, and the one associated only with CIII_2_ as CIV_C_. A composite map of SC and the locally refined maps of three CIV unis are shown in the Figures 4A and 4B, respectively.

### Membrane Curvature Induced by SCs

Structural comparison of revealed a modest curvature of the membrane for the CI_1_CIII_2_ in the presence of CIV_A_ obtained from the previous porcine mitochondria study compared to the CI_1_CIII_2_ SC (Figure S2B). In contrast, comparison of our CI_1_CIII_2_CIV_3_ SCs map derived from SMPs with the previously reported map of the CI_1_CIII_2_CIV_1_ SCs from intact porcine mitochondria showed nearly identical membrane curvature across all viewing angles (Figure S2C). These observations suggest that the membrane curvature observed in CI_1_CIII_2_CIV SCs is primarily attributable to the incorporation of CIV_A_.

### Megacomplexes in SMPs membranes

Previous studies have identified a larger SCs unit, termed the respiratory megacomplex, where CI with CIV associates with CI_1_CIII_2_CIV_1_^26^ to form a circular complex, CI_2_CIII_2_CIV_2_^27^ (Figure S3D). Similar respiratory megacomplexes have also been reported on the mitochondrial inner membrane^21,28^.

To investigate whether such mega-complexes are present in SMP membranes, we re-extracted 764,000 particles—predominantly containing SCs—using an expanded box size of 981 Å. Subsequent 2D classification yielded class averages revealed chromosome-shaped SCs (Figure S3A). This 2D projection closely resembled the previously reported CI_2_CIII_2_CIV_2_ SCs identified in porcine heart mitochondria; however, four additional densities—putatively corresponding to CIV—were clearly discernible in our data, suggesting that the observed structure is more likely to represent CI_2_CIII_2_CIV_6_ (Figure S3A). In this study, the predominant SCs identified on the SMPs were of the CI_1_CIII_2_CIV_3_ type. In contrast, CI_2_CIII_2_CIV_2_ type SCs, in which two CI units associate with a single CIII_2_ (Figure S3D), were not observed on the SMPs. This absence may be due to the presence of CIV_B_ and CIV_C_ in the CI_1_CIII_2_CIV_3_ SCs, which likely sterically hinder the association of an additional CI with CIII_2_. These findings suggest that SCs with diverse subunit compositions, including higher-order assemblies, are embedded within the SMP membranes.

Additionally, 2D class averages revealed a mushroom-shaped density adjacent to the SC (Figure S3B, indicated by red arrow). A 3D reconstruction of this particle class unambiguously resolved a structure resembling the monomeric F_1_ head (Figure S3B, right), implying that F_o_F_1_ ATP synthase is distributed not only in regions of membrane curvature but also in more planar regions of the membrane.

### Structures of complex I in SCs

Complex I (CI) is the largest protein complex in the mitochondrial respiratory system, comprising 45 distinct subunits with a total molecular mass of approximately 1 MDa^5^. It adopts a characteristic L-shaped architecture, consisting of a hydrophilic domain responsible for electron transfer and a membrane-embedded domain responsible for proton translocation (Figure 5A and 5B). The ubiquinone (UQ) substrate binds within a tunnel-like structure located at the interface between these two domains, known as the “UQ-accessing tunnel”, which is formed by the 49-kDa, PSST, ND1, ND6, and ND3 subunits.

**Figure 5.**
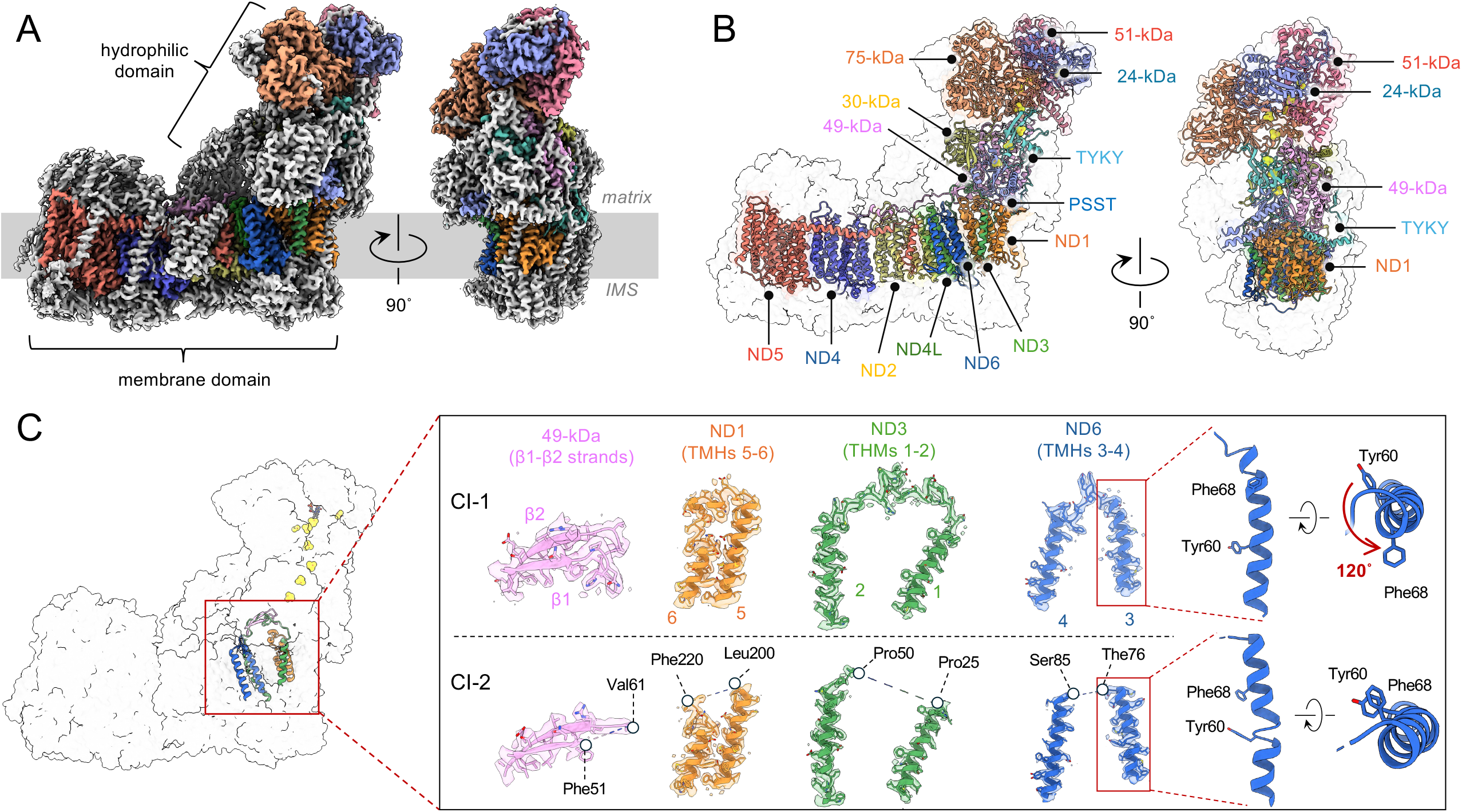
Structure of complex I (CI-1) in the supercomplex (SC) and conformational differences between CI-1 and CI-2. (A) Cryo-EM density map of CI-1. The density corresponding to the 14 core subunits is color-coded. (B) Atomic model of CI-1 built from the corresponding density map. The 14 core subunits are shown in color, while the 30 supernumerary (accessory) subunits are rendered in gray. (C) Comparison of subunits exhibiting density differences between CI-1 and CI-2. Overlaid densities and atomic models for each class are presented. In CI-2, certain regions of the 49-kDa, ND1, ND6, and ND3 subunits show poorly resolved densities, precluding accurate model building. Adjacent amino acid residues are annotated near regions where density is absent in CI-2.

Classification focused on complex I (CI) within the SC (CI_1_CIII_2_CIV_3_) revealed two distinct subclasses, termed CI-1 and CI-2 at a resolution of 2.55 and 2.57-Å, respectively. CI-1 displayed a well-ordered architecture at the interface between the hydrophilic and membrane-embedded domains, whereas CI-2 adopted a more disordered conformation. CI-2 showed disorder in four loop regions: the loop connecting the β1–β2 strands of the 49-kDa subunit, that between transmembrane helices (TMHs) 5–6 of ND1, that between TMHs 4–5 of ND6, and that between TMHs 1–2 of ND3 (Figure 5C). These structural hallmarks align with the closed and open conformations of CI, which have been extensively described and characterized by multiple research groups, where the putative UQ-accessing tunnel adopts either a relaxed or constrained state, respectively.

In the CI-1 (closed) conformation, a density was detected within the UQ-accessing tunnel (Figure S4A). Although a UQ_10_ molecule could be fitted into this density without significant steric clashes (Figure S4B), the local resolution is insufficient to confidently model UQ_10_. Therefore, we do not assign this density to UQ_10_, and its molecular identity remains uncertain.

Overall, the CI-1 and CI-2 structures closely resemble the active- and deactive-conformations of CI observed *in situ* in porcine heart mitochondria (PDB IDs: 8UEO and 8UES), where the UQ-access cavities adopt closed- and open-conformations, respectively^5^. The RMSDs relative to these structures are approximately 1.11 Å for CI-1 and 1.13 Å for CI-2. Furthermore, CI-1 is nearly identical to the closed-state structure of isolated bovine heart CI (PDB ID: 7QSK)^29^, with an RMSD of ∼0.89 Å. The close structural similarity between our cryo-EM structures and those previously reported by other groups—derived from both *in situ* and isolated preparations—suggests that CI retains a high degree of structural integrity even during biochemical procedures, including membrane solubilization and sonication, highlighting its intrinsic robustness.

### Structures of complex III dimer in SCs

Within the SCs (CI_1_CIII_2_CIV_3_) of SMPs, complex III (CIII) exists as a two-fold symmetric dimer (CIII_2_), with each monomer comprising 11 distinct subunits (Figures 6A and 6B). CIII_2_ associates with CI through multiple contacts: CI-B15 (NDUFB4) and CI-B22 (NDUFB9) with CIII–core protein 1 on the matrix side, and CI-B14.7 (NDUFA11) with CIII-subunit 8 within the membrane bilayer (Figure S5A).

**Figure 6.**
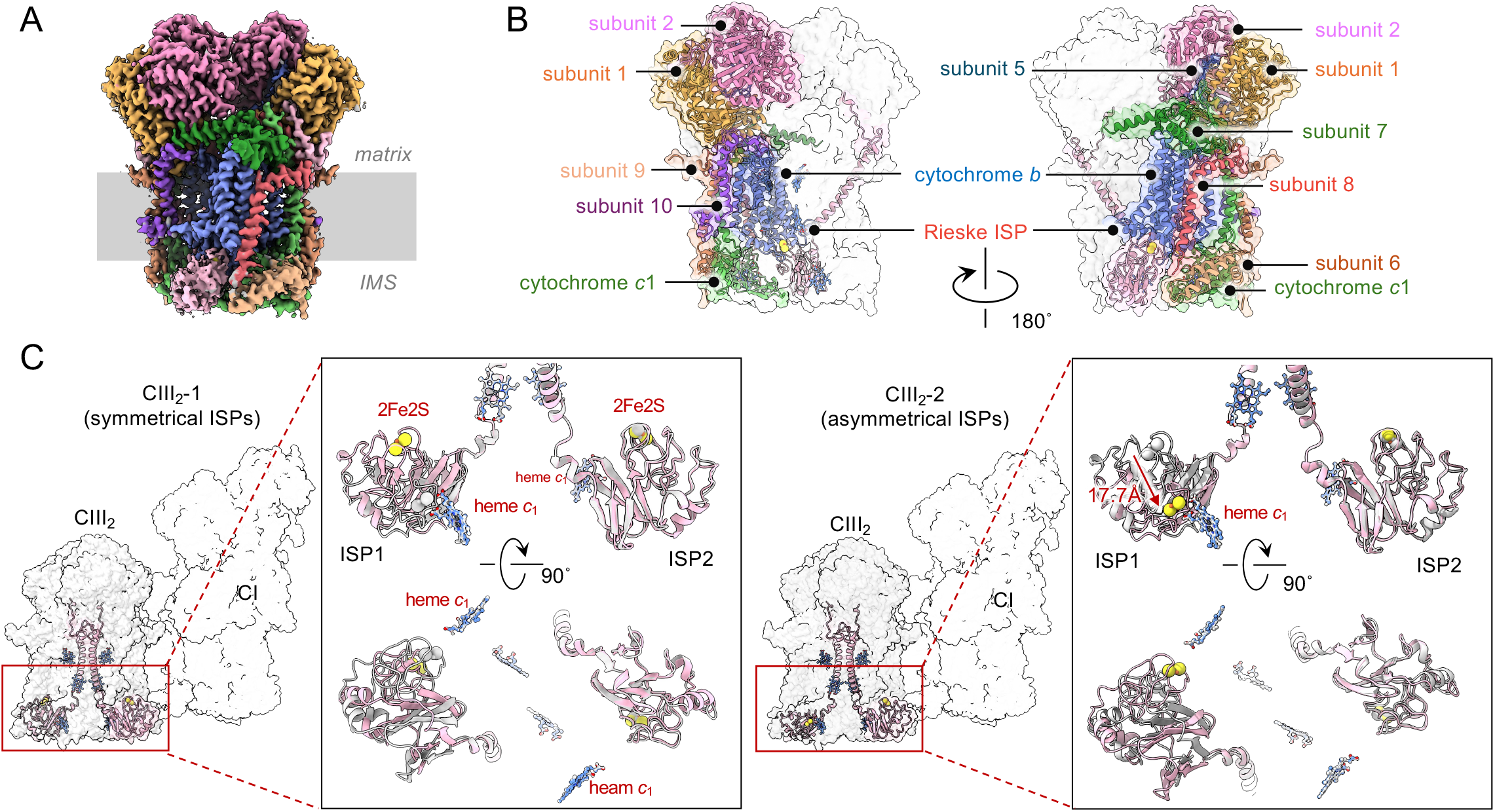
Structure of CIII_2_ in the supercomplex (SC) and conformational differences in the ISP domain between CIII_2_-1 and CIII_2_-2. (A) Cryo-EM density map of CIII_2_-1. The density corresponding to each subunit is shown in a different color. (B) Atomic model of CIII_2_-1, with subunits color-coded as in panel (A). (C) Structural comparison of CIII_2_-1 and CIII_2_-2. The overall views show the relative positioning of CIII_2_ and CI within the SC. Insets present enlarged views of the ISP head domains, where CIII_2_-1 (light pink) is overlaid with CIII_2_-2 (gray, left panel), and vice versa (right panel). These overlays reveal a marked conformational difference in ISP1: in CIII_2_-2, the 2Fe– 2S cluster of ISP1 is repositioned 17.7 Å closer to the heme c_1_ of cytochrome c_1_ compared to its position in CIII_2_-1.

Focused classification of the cryo-EM data resolved CIII_2_ into two structural classes, CIII_2_-1 and CIII_2_-2, at resolutions of 2.23 Å and 2.42 Å, respectively (Figure S1C). For both classes, complete atomic models were built, including all amino acid residues and cofactors. The overall architectures of CIII_2_-1 and CIII_2_-2 were nearly identical, except for the conformation of the iron–sulfur protein (ISP), a core catalytic subunit containing a 2Fe–2S cluster responsible for electron transfer from heme *b*_H_ of cytochrome *b* to heme *c*_1_ of cytochrome *c*_1_. In CIII_2_-1, the ISPs adopt symmetric conformations, whereas in CIII_2_-2, they exhibit asymmetry (Figure 6C). Specifically, in CIII_2_-2, one solvent-exposed Rieske domain—the mobile part of the ISP—remains in the same position as in CIII_2_-1, while the other shifts toward heme c_1_ (Figure 6C).

Previous cryo-EM studies by Wieferig and Kühlbrandt on CIII_2_ isolated from *Yarrowia lipolytica* under varying redox conditions demonstrated that the Rieske domain adopts multiple conformations to facilitate electron transfer between hemes *b*_L_ and *c*_1_ via the 2Fe–2S cluster^30^. They also reported both symmetric and asymmetric CIII_2_ dimers, depending on the relative positions of the two Rieske domains. Although it remains debated whether the two ISPs operate independently or in a coordinated manner, our structures of bovine CIII_2_ replicate the asymmetric conformation observed in the *Yarrowia* enzyme, suggesting that each Rieske domain moves independently, potentially reflecting the redox state of each monomer.

### Structures of complex IV in SCs

Among the three complex IV (CIV) units present within the SC (CI_1_CIII_2_CIV_3_), local refinement of CIV_A_—positioned adjacent to the distal end of complex I via interactions between the CI-ND5 and CIV_A_-Cox7A subunits (Figure S5B)—yielded a density map at 2.61 Å resolution, enabling atomic model construction of CIV_A_ (Figure 7A). Consistent with recent findings, including the *in situ* cryo-EM structure of porcine mitochondrial CIV^16^, an additional density corresponding to the 14th subunit, NDUFA4, was identified in all three CIV units (Figures 7B).

**Figure 7.**
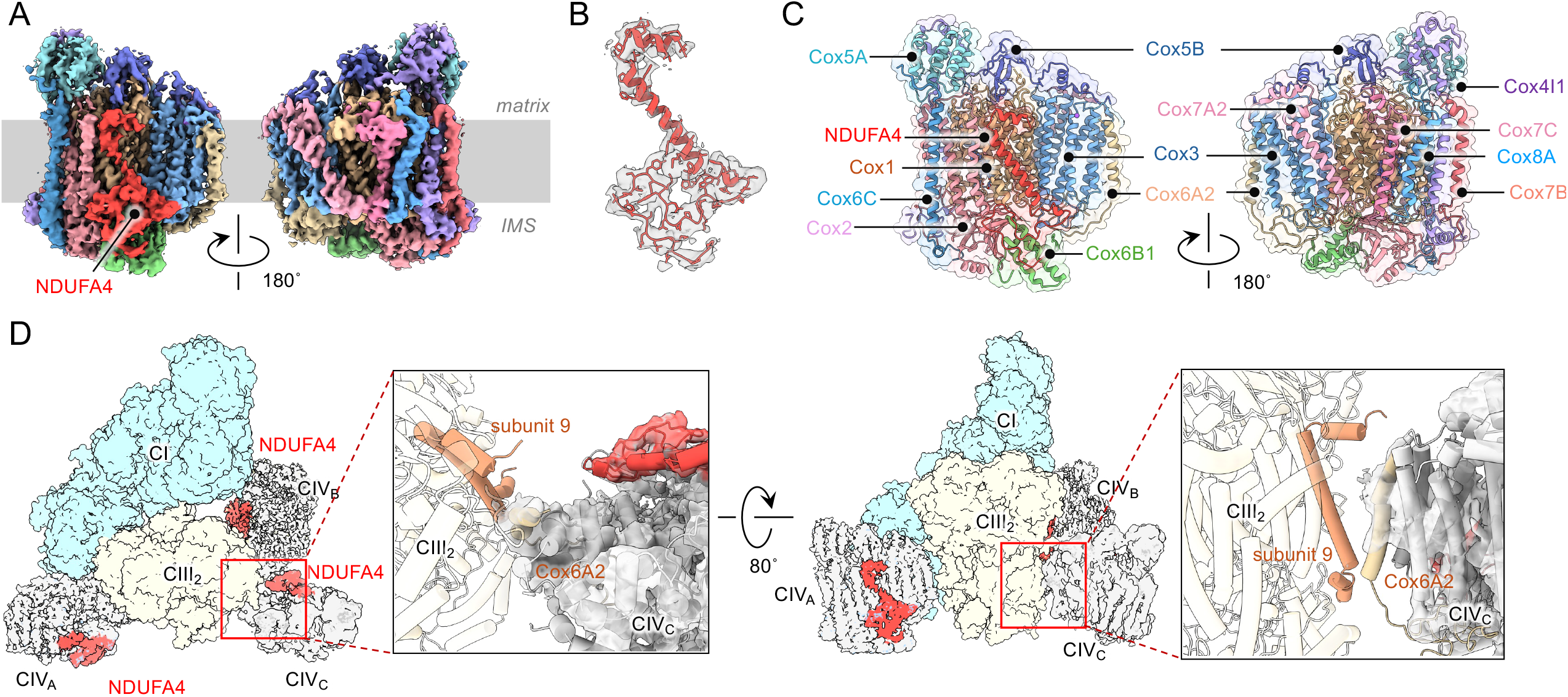
Structure of CIV_A_ in the supercomplex (SC) and interaction between CIV_C_ and CIII_2_. (A) Cryo-EM density map of CIV_A_. Each subunit is color-coded. (B) Close-up view of the newly identified NDUFA4 subunit. The atomic model is superimposed on the corresponding EM density (shown in gray). (C) Atomic model of CIV_A_, with subunits color-coded as in panel (A). (D) Interface between CIV_C_ and CIII_2_ within the SC. The left panel shows a top-down view of the interface between CIII_2_ and CIV_C_, while the right panel presents a side view. The model of CIV_C_ (shown in a tube helix representation) is superimposed on the corresponding EM density map. To clarify the orientation of CIV_C_ within the SC, the subunit NDUFA5 is highlighted in red.

The second CIV unit, CIV_B_ (2.86 Å resolution), was located in the interspace between CI and CIII_2_, forming contacts with both complexes—specifically between CI-39-kDa and CIV_B_-Cox5A on the matrix side, and between CIII-subunit 9 and CIV_B_-Coq6A2 within the membrane bilayer (Figure S5C). The third CIV unit, CIV_C_ (3.98 Å resolution), was positioned adjacent to CIII_2_. Although its resolution was lower than that of CIV_A_ and CIV_B_, the TMHs remained clearly discernible, allowing confident identification (Figure S1C). At the interface between CIV_C_ and CIII_2_, the TMH of CIV_C_ -Cox6A2 was located in close proximity to CIII_2_-subunit 9 (Figures 7C). However, due to the limited resolution of CIV_C_, it was not possible to observe how it associates with CIII_2_ at the amino acid level—whether through direct protein–protein interactions or via lipid-mediated association.

## Discussion

### F_o_F_1_ oligomers in SMPs membrane

In this study, we successfully determined the structures of oligomeric F_o_F_1_ ATP synthase and respiratory SCs located in the mitochondrial inner membrane without the need for extraction and isolation using detergent. We determined the oligomeric structure of F_o_F_1_ on SMPs, revealing the existence of dimers, in which two F_o_F_1_ monomers are linked by dimeric IF_1_ (Figure 2). Using focused refinement on each monomer, we obtained the structure of monomeric F_o_F_1_ at a resolution sufficient to construct an atomic model. Furthermore, we identified a tetrameric structure with an additional F_o_F_1_-IF_1_ dimer opposite side to the originally identified F_o_F_1_-IF_1_ dimer (Figure 3A).

In the tetrameric F_o_F_1_ structure obtained in this study, the F_o_F_1_-IF_1_ dimer pairs are associated through their membrane domains, inducing a steep membrane curvature (Figure 3A, lower). Additionally, density corresponding to the F_1_ heads was observed on both sides of the reconstructed F_o_F_1_ tetramer (Figure 3A, right). These findings suggest that the F_o_F_1_ tetramer structures align laterally, indicating the presence of a curved structure on the SMPs membrane similar to the cristae tips in mammalian mitochondria. The involvement of the F_o_F_1_ tetramer structure in membrane curvature has been previously suggested by detergent solubilized F_o_F_1_ tetramer structures, and our results provide direct evidence supporting this hypothesis, in a mammalian biological membrane. Our analysis also revealed a multimeric structure in which the F_o_F_1_-IF_1_ dimer was offset by one F_o_F_1_ unit (Figure 3B), suggesting association of F_o_F_1_-IF_1_ dimers through the F_o_ domain was weak at cristae tips. In ciliates, a stable F_o_F_1_ dimer structure formed via interactions between the peripheral stalks and F_o_ domains has been reported^30^, and this dimer configuration is known to contribute to cristae tip formation in the mitochondria of these organisms. In contrast, here the interaction between the F_o_F_1_ monomers at the tip of curved structure in the F_o_F_1_ tetramer appears to rely solely on the F_o_ domain. Our observation of the offset F_o_F_1_-IF_1_ dimer structure implys that the architecture of cristae tips is inherently dynamic. Indeed, cristae morphology has been shown to undergo dramatic changes depending on the metabolic state of mammalian mitochondria, and the fluidity of F_o_F_1_ multimeric assemblies may play a role in facilitating such structural remodeling^31^.

In addition, we have also obtained cryoEM images showing the presence of a monomeric F_1_ heads near the SCs (Figure S3). While most F_o_F_1_ complexes are located at the tips of cristae-like membrane structures, the presence of such orphan F_o_F_1_ is suggestive of the fluidity of mitochondrial cristae structures.

### Structural organization of respiratory supercomplexes

We also successfully obtained a high-resolution 3D structure of the respiratory SCs directly from SMPs. The predominant SC exhibited a stoichiometry of CI_1_CIII_2_CIV_3_, in which three CIV units are bound to the CI_1_CIII_2_ core. This organization differs from SCs derived from solubilized mitochondrial membrane, which typically contain only a single CIV unit, as well as from the *in situ* structure of CI_1_CIII_2_CIV_2_ observed in porcine heart mitochondria, where the second CIV is positioned between the ND6 subunit of CI and the cytochrome *b* subunit of CIII_2_^16^. Moreover, 2D class averages revealed a novel megacomplex with the stoichiometry CI_2_CIII_4_CV_6_ (Figure S3A). In contrast, the structure corresponding to CI_2_CIII_2_CV_2_—previously reported in *in situ* structural analyses of porcine heart mitochondria^16^—were not identified. In the CI_2_CIII_2_CV_2_ complex (Figure S3D), the two CI units are bridged by a shared CIII_2_; therefore, binding of CIV to CIII_2_ may sterically hinder the formation of this arrangement.

As noted in previous studies of porcine mitochondrial SCs^16^, solubilization of membrane proteins using detergents may disrupt SC architecture by simultaneously stripping off tightly bound phospholipids that are essential for stabilizing interactions among respiratory complexes. In this study, structural determination was achieved without solubilizing the mitochondrial inner membrane, enabling the identification of novel SCs, CI_1_CIII_2_CIV_3_ and CI_2_CIII_4_CV_6_. This detergent-free approach may facilitate the discovery of previously unidentified SCs, not only within the mitochondrial inner membrane but also across other organellar or plasma membranes.

### Structures of ubiquinone-accessing tunnel in Complex I

Compared to the F_o_F_1_ oligomer, the respiratory SCs were resolved at higher resolution, enabling the construction of atomic models for each respiratory complex. Within the SCs, complex I (CI) was successfully classified into closed and open conformations (Figure 5C and Figure S4), consistent with those previously reported for the isolated enzyme. Notably, these conformations likely correspond to the biochemically defined active and deactive states of CI, respectively; however, this correspondence remains a subject of ongoing debate^5, 6^.

In the closed conformation structure, the narrow ubiquinone (UQ)-accessing tunnel is occupied by a density likely corresponding to UQ (Figure S4), which appears to hinder the binding of additional ligands. Nevertheless, our chemical biology studies using bovine SMPs revealed that bulky synthetic ligands—larger than the tunnel diameter in the closed conformation—can still access the UQ-binding sites^32,33^. This observation suggests that the open conformation, whose complete structure remains unresolved, permits access for such ligands. These findings support the possibility that the open conformation of CI is competent for ligand binding. Further investigations integrating structural analyses of SCs in bovine SMPs with our chemical biology approaches provide insights into the relationship between closed/open conformations and the active/inactive states of CI, thereby advancing our understanding of the mechanisms underlying their structural transitions.

### Perspective

Recent studies have reported that metabolic processes are regulated through alterations in the composition of mitochondrial respiratory SCs^28^. The isolation of mitochondria from small biopsy specimens, particularly from patients with mitochondrial disorders or metabolic diseases such as diabetes, holds promise for uncovering structural correlations between SCs and disease pathophysiology. Notably, given that cyt *c*_*1*_ is released into the cytosol during the early stages of apoptosis, *in situ* structural analysis of SCs and F_o_F_1_ oligomer from apoptotic cells may provide critical insights into the molecular mechanisms governing initiation of apoptosis. Collectively, these advances underscore the potential of native-state structural biology to deepen our understanding of mitochondrial function and its implications in human health and disease.

## Acknowledgements

We are grateful to all the members of the Yokoyama Lab for their continuous support and technical assistance. We are thankful to Dr. Jun-ichi Kishikawa (Kyoto Institute of Technology) for the structural analysis. Our research was supported by Grant-in-Aid for Scientific Research (JSPS KAKENHI) Grant Numbers 23H02453 to K.Y., 24K08729 to T.M. and 25K01958 to M.M., Takeda Science foundation funding to K.Y., The Uehara Memorial foundation to M.M, and Tokyo Kasei Chemical Promotion foundation to M.M. Our research was also supported by the Platform Project for Supporting Drug Discovery and Life Science Research (Basis for Supporting Innovative Drug Discovery and Life Science Research (BINDS)) from AMED under Grant Number JP17am0101001 (support number 1312), and Grants-in-Aid from the “ARIMS” of the Ministry of Education, Culture, Sports, Science and Technology (MEXT) to KY.

## Author contributions

A.N., and K.Y. designed, performed and analyzed the experiments. A.N., T.M., S.A., M.I., analyzed the data and contributed to preparation of the samples. K.M. provided technical support and conceptual advice. A.N., H.M., M.M., K.Y. designed and supervised the experiments and wrote the manuscript. All authors discussed the results and commented on the manuscript. The authors declare no conflicts of interest associated with this manuscript. All data is available in the manuscript or in the supplementary materials.

## Data and materials availability

The Cryo-EM density maps and models generated in this study have been deposited in the EM and protein database under accession codes: EMDB 65580, EMDB 65581, EMDB 65583, EMDB 65584, EMDB 65585, EMDB 65586, EMDB 65587, EMDB 65577, EMDB 65578, EMDB 65579, PDB 9W2U, PDB 9W2V, PDB 9W2X, PDB 9W2Y,

PDB 9W2Z, PDB 9W2R, PDB 9W2S, PDB 9W2T. The deposited PDB files, cryo-EM density maps, and validation reports have been uploaded to Figshare.

## Material and Methods

### Preparation of SMPs

Bovine heart mitochondria were isolated according to the method described by Smith^34^. Heart muscle from one bovine heart, obtained from slaughterhouse within several hours post-slaughter, was minced using a meat grinder at 4 °C to yield approximately 1 kg of tissue. The mince was disrupted using a mechanical blender in a buffer containing 250 mM sucrose, 1.0 mM sodium succinate, 0.2 mM EDTA, and 10 mM Tris-HCl (pH 7.8) at a ratio of 300 g mince/L of buffer. The pH was subsequently adjusted to 7.4 with 6.0 M KOH. The homogenate was centrifuged at 680 × g for 20 min at 4 °C to remove unbroken cells and nuclei. The resulting supernatant was neutralized to pH 7.4 with 1.0 M KOH and centrifuged at 10,000 × *g* for 20 min at 4°C. The pellet was carefully homogenized using a tight-fitting Teflon-glass homogenizer (100 ml capacity) and centrifuged at 30,700 × *g* for 15 min at 4°C. This homogenization and centrifugation step was repeated, and the final pellet containing both light- and heavy-mitochondria was resuspended in the same buffer (∼50 ml) and stored at −80 °C as intact mitochondria until use. The protein concentration was determined using the Biuret method. Typically, 1,500-2,000 mg of mitochondria were obtained from one bovine heart.

Submitochondrial particles (SMPs) were prepared as described by Matsuno-Yagi and Hatefi^24^. Frozen mitochondria were thawed and adjusted to a final protein concentration of 40 mg of protein/ml with a buffer containing 250 mM sucrose and 10 mM Tris/HCl (pH 7.5). The suspension was supplemented with 1.0 mM potassium succinate, 1.5 mM ATP, 10 mM MgCl_2_, 10 mM MnCl_2_, followed by a sonication using a TOMY UR-200P (power setting 8, 30 sec, 5 cycles, with 3 min intervals) on ice. After sonication, the pH was adjusted to 7.5 with 1.0 M KOH and the sample centrifuged for 15 min at 34,400 x *g* at 4°C. The clear supernatant was carefully collected and subjected to ultracentrifugation at 200,000 x *g* for 45 min at 4°C. The resulting brown pellet was rinsed with a buffer containing 250 mM sucrose and 10 mM Tris/HCl (pH 7.5) and homogenized in the same buffer at a protein concentration of 30-40 mg/ml. The SMP suspension was snap-frozen in liquid nitrogen in small aliquots (100 µL) and stored at −80°C.

### Cryo-grid preparation and image acquisition for SMPs

SMP suspensions (8 mg/ml) were used for preparation of cryogrids using Quantifoil UltraAufoil R1.2/1.3 grids. Before blotting, the grids were treated with an ion bombardier for 1 minute of glow discharge. Using a FEI Vitrobot (Thermo Fisher Scientific), 3 µl of SMP solution were applied to a grid using a blot time of 2.5 s, blot force 10, and a drain time of 0.5 s at 25°C and 100% humidity. Cryo-grids were imaged using a Titan Krios G2 (Thermo Fisher Scientific) (UHV EM at Osaka University) equipped with a K3 electron detector (Gatan). For data collection using the Titan Krios G2 (Thermo Fisher Scientific), cryo-EM movies were collected automatically at a nominal magnification of 29,000x corresponding to a calibrated pixel size of 0.84 Å/pix using image shift based data collection with the SerialEM software employing a defocus range of −0.8 to −1.8 μm. Image data was collected in 50 frames in CDS mode at a total electron dose of 50 electrons/Å^2^.

### Structural analysis

The detailed single particle analysis workflow is summarized in Figure S1. Single particle analysis was performed by RELION ver 4.0^35^ and CryoSPARC ver 4.7^36^. Particle metadata in cryoSPARC (.cs) format were converted to RELION .star files using the PyEM script csparc2star.py. Beam-induced drift motion was corrected in 30,680 movies using MotionCor2^37^, CTF parameters were estimated with CTFFIND4^38^. Approximately 300 particles of F_o_F_1_ dimer or SC were manually picked and used to train Topaz. Using the trained model, 3,636,680 particles were picked from 5,000 micrographs, extracted with a box size of 150 pixels (4.36 Å/pix), and 2D classification was performed. Particles of F_o_F_1_ dimer or SCs were selected and initial maps of F_o_F_1_ dimer and SCs were created by Ab-initio reconstruction, respectively. Heterogeneous Refinements were then performed with all the particles picked by topaz to obtain F_o_F_1_ dimer or SCs structures. The topaz models were further trained using 3000 particles of the both F_o_F_1_ dimer and SCs particles curated by Heterogeneous Refinement. Using both models, a total of 40,298,925 particles were picked and extracted from 30,680 micrographs. These particles were divided into eight subsets, with obvious junk particles excluded by 2D classification, yielding 30,421,492 particles.

Heterogeneous refinement was performed four times using either the F_o_F_1_ dimer or SCs, and multiple junk classes resulting from Ab-initio reconstruction as reference volumes. To obtain high-precision CTF parameters, the 764,363 particles of the SCs were processed with C2 symmetry imposed on the Complex III region, followed by Local refinement and combined Global/Local CTF refinement. Iteration of this process yielded a structure of 2.3 Å resolution for the CIII_2_ domain. The particles with CTF parameters refined in the CIII_2_ region were then used to proceed with structural analysis on the other respiratory chain proteins. After local refinement in the CI, 3D classification was performed to classify the CI as open or closed, and the respective structures were obtained. For the CIII_2_, symmetry expansion was performed on C2, and after local refinement, 3D classification was performed using a mask including the Rieske domain to classify the CIII_2_-1 and CIII_2_-2. For CIV_A_, CI_1_CIII_2_ and CI_1_CIII_2_CIV_A_ each was classified by Heterogeneous Refinement, followed by Local Refinement and then 3D classification to remove low-resolution classes to obtain the structure. For CIV_B_ and CIV_C_, Heterogeneous Refinement classified structures with stronger densities of CIV_B_ and CIV_C_. Then, CI_1_CIII_2_CIV_A_ domain were signal subtracted to improve alignment in Local Refinement. The structures of CIV_B_ and CIV_C_ were obtained by removing the junk class by 3D classification.

For structural analysis of the megacomplex and SCs-F_o_F_1_, we re-extracted at a 981pixel box size (3.27 Å/pix) and limited the alignment resolution to 20-30 Å in the 2D classification and clear 2D class-averaged images of both the megacomplex and SC-F_o_F_1_ were obtained. By performing Heterogeneous Refinement of these particles with the structure of SCs as a reference, followed by Homogeneous refinement, the 3D structures of the megacomplex and SC-F_o_F_1_ were obtained. For analysis of the F_o_F_1_ oligomer, Heterogeneous Refinement yielded 328,822 particles, and the structure of the F_o_F_1_-IF_1_ dimer was obtained by Homogeneous Refinement. First, CTF refinement and Local refinement were performed on the F_1_-1 domain, followed by local refinement and 3D classification in F_o_-1, F_1_-2, and F_o_-2 to obtain the structure of each. To obtain structural information of a broader region of the F_o_F_1_ oligomer, particles were re-extracted using a box size of 981 pixels (3.27 Å/pixel), followed by local refinement and 3D classification focused on the region encompassing the F_o_F_1_–IF_1_ dimer and adjacent F_o_F_1_ units. Through 3D classification, a distinct class was identified corresponding to a tetrameric assembly in which two F_o_F_1_–IF_1_ dimers are paired. Using this structure as a reference, heterogeneous refinement further resolved two conformational states of the F_o_F_1_ tetramer.

### Model building

The atomic model of F_o_F_1_ monomer and focused F_o_ domain were built from the Cryo-EM structure of bovine ATP synthase PDB 6ZPO, 6ZQN, and 6ZBB. For the model construction of SCs, we used bovine CI structure of PDB7QSK, bovine CIII structure of 1SQB, and CIV structure of 6JY3. Rigid body fitting was performed by ChimeraX^39^, and after manual modification of the entire model by ISOLDE^40^, a refinement by phenix.real_space_refinement^41^ was performed. The refinement model was evaluated with MolProbity^42^, and Phenix.real_space_refinement. Manual correction with COOT and ISOLDE and phenix.real_space_refinement were repeated until model parameters improved.

## References

1. Levental, I. & Lyman, E. Regulation of membrane protein structure and function by their lipid nano-environment. Nat Rev Mol Cell Biol 24, 107–122 (2023).

2. Payandeh, J. & Volgraf, M. Ligand binding at the protein-lipid interface: strategic considerations for drug design. Nat Rev Drug Discov 20, 710–722 (2021).

3. Kuhlbrandt, W. Structure and Mechanisms of F-Type ATP Synthases. Annu Rev Biochem 88, 515–549 (2019).

4. Letts, J.A., Fiedorczuk, K., Degliesposti, G., Skehel, M. & Sazanov, L.A. Structures of Respiratory Supercomplex I+III(2) Reveal Functional and Conformational Crosstalk. Mol Cell 75, 1131–1146.e6 (2019).

5. Hirst, J. Mitochondrial complex I. Annu Rev Biochem 82, 551–75 (2013).

6. Sazanov, L.A. A giant molecular proton pump: structure and mechanism of respiratory complex I. Nat Rev Mol Cell Biol 16, 375–88 (2015).

7. Sarewicz, M. et al. Catalytic Reactions and Energy Conservation in the Cytochrome bc(1) and b(6)f Complexes of Energy-Transducing Membranes. Chem Rev 121, 2020–2108 (2021).

8. Cecchini, G. Function and structure of complex II of the respiratory chain. Annu Rev Biochem 72, 77–109 (2003).

9. Berry, E.A., Guergova-Kuras, M., Huang, L.S. & Crofts, A.R. Structure and function of cytochrome bc complexes. Annu Rev Biochem 69, 1005–75 (2000).

10. Kaila, V.R., Verkhovsky, M.I. & Wikström, M. Proton-coupled electron transfer in cytochrome oxidase. Chem Rev 110, 7062–81 (2010).

11. Shimada, A., Tsukihara, T. & Yoshikawa, S. Recent progress in experimental studies on the catalytic mechanism of cytochrome c oxidase. Front Chem 11, 1108190 (2023).

12. Vinothkumar, K.R., Zhu, J. & Hirst, J. Architecture of mammalian respiratory complex I. Nature 515, 80–84 (2014).

13. Sun, F. et al. Crystal structure of mitochondrial respiratory membrane protein complex II. Cell 121, 1043–57 (2005).

14. Iwata, S. et al. Complete structure of the 11-subunit bovine mitochondrial cytochrome bc1 complex. Science 281, 64–71 (1998).

15. Tsukihara, T. et al. Structures of metal sites of oxidized bovine heart cytochrome c oxidase at 2.8 A. Science 269, 1069–74 (1995).

16. Zheng, W., Chai, P., Zhu, J. & Zhang, K. High-resolution in situ structures of mammalian respiratory supercomplexes. Nature 631, 232–239 (2024).

17. Yokoyama, K. Rotary mechanism of V/A-ATPases-how is ATP hydrolysis converted into a mechanical step rotation in rotary ATPases? Front Mol Biosci 10, 1176114 (2023).

18. Boyer, P.D. The binding change mechanism for ATP synthase--some probabilities and possibilities. Biochim Biophys Acta 1140, 215–50 (1993).

19. Zhou, A. et al. Structure and conformational states of the bovine mitochondrial ATP synthase by cryo-EM. Elife 4, e10180 (2015).

20. Spikes, T.E., Montgomery, M.G. & Walker, J.E. Structure of the dimeric ATP synthase from bovine mitochondria. Proc Natl Acad Sci U S A 117, 23519–23526 (2020).

21. Gu, J. et al. Cryo-EM structure of the mammalian ATP synthase tetramer bound with inhibitory protein IF1. Science 364, 1068–1075 (2019).

22. Gerle, C. et al. Human F-ATP synthase as a drug target. Pharmacol Res 209, 107423 (2024).

23. Chinnery, P.F. Primary Mitochondrial Disorders Overview. in GeneReviews(®) (eds. Adam, M.P. et al.) (University of Washington, Seattle Copyright © 1993-2025, University of Washington, Seattle. GeneReviews is a registered trademark of the University of Washington, Seattle. All rights reserved., Seattle (WA), 1993).

24. Matsuno-Yagi, A. & Hatefi, Y. Studies on the mechanism of oxidative phosphorylation. Flow-force relationships in mitochondrial energy-linked reactions. J Biol Chem 262, 14158–63 (1987).

25. Letts, J.A., Fiedorczuk, K. & Sazanov, L.A. The architecture of respiratory supercomplexes. Nature 537, 644–648 (2016).

26. Wu, M., Gu, J., Guo, R., Huang, Y. & Yang, M. Structure of Mammalian Respiratory Supercomplex I(1)III(2)IV(1). Cell 167, 1598–1609.e10 (2016).

27. Guo, R., Zong, S., Wu, M., Gu, J. & Yang, M. Architecture of Human Mitochondrial Respiratory Megacomplex I(2)III(2)IV(2). Cell 170, 1247–1257.e12 (2017).

28. Liang, C. et al. Formation of I(2)+III(2) supercomplex rescues respiratory chain defects. Cell Metab 37, 441–459.e11 (2025).

29. Chung, I. et al. Cryo-EM structures define ubiquinone-10 binding to mitochondrial complex I and conformational transitions accompanying Q-site occupancy. Nat Commun 13, 2758 (2022).

30. Dietrich, L., Agip, A.A., Kunz, C., Schwarz, A. & Kühlbrandt, W. In situ structure and rotary states of mitochondrial ATP synthase in whole Polytomella cells. Science 385, 1086–1090 (2024).

31. Kondadi, A.K., Anand, R. & Reichert, A.S. Cristae Membrane Dynamics - A Paradigm Change. Trends Cell Biol 30, 923–936 (2020).

32. Masuya, T., Murai, M., Ifuku, K., Morisaka, H. & Miyoshi, H. Site-specific chemical labeling of mitochondrial respiratory complex I through ligand-directed tosylate chemistry. Biochemistry 53, 2307–17 (2014).

33. Masuya, T. et al. Pinpoint Chemical Modification of the Quinone-Access Channel of Mitochondrial Complex I via a Two-Step Conjugation Reaction. Biochemistry 56, 4279–4287 (2017).

34. Smith, A.L. Preparation, properties, and conditions for assay of mitochondria: Slaughterhouse material, small-scale. Methods Enzymol 10, 81–86 (1967).

35. Kimanius, D., Dong, L., Sharov, G., Nakane, T. & Scheres, S.H.W. New tools for automated cryo-EM single-particle analysis in RELION-4.0. Biochem J 478, 4169–4185 (2021).

36. Punjani, A., Rubinstein, J.L., Fleet, D.J. & Brubaker, M.A. cryoSPARC: algorithms for rapid unsupervised cryo-EM structure determination. Nat Methods 14, 290–296 (2017).

37. Zheng, S.Q. et al. MotionCor2: anisotropic correction of beam-induced motion for improved cryo-electron microscopy. Nat Methods 14, 331–332 (2017).

38. Rohou, A. & Grigorieff, N. CTFFIND4: Fast and accurate defocus estimation from electron micrographs. J Struct Biol 192, 216–21 (2015).

39. Pettersen, E.F. et al. UCSF ChimeraX: Structure visualization for researchers, educators, and developers. Protein Sci 30, 70–82 (2021).

40. Croll, T.I. ISOLDE: a physically realistic environment for model building into low-resolution electron-density maps. Acta Crystallogr D Struct Biol 74, 519–530 (2018).

41. Afonine, P.V. et al. Real-space refinement in PHENIX for cryo-EM and crystallography. Acta Crystallogr D Struct Biol 74, 531–544 (2018).

42. Chen, V.B. et al. MolProbity: all-atom structure validation for macromolecular crystallography. Acta Crystallogr D Biol Crystallogr 66, 12–21 (2010).

