## Supplementary materials for "*In situ* structure determination of Respiratory Supercomplexes and ATP synthase oligomers in mammalian mitochondrial inner membrane"

Figures S1-5

Tables S1-2

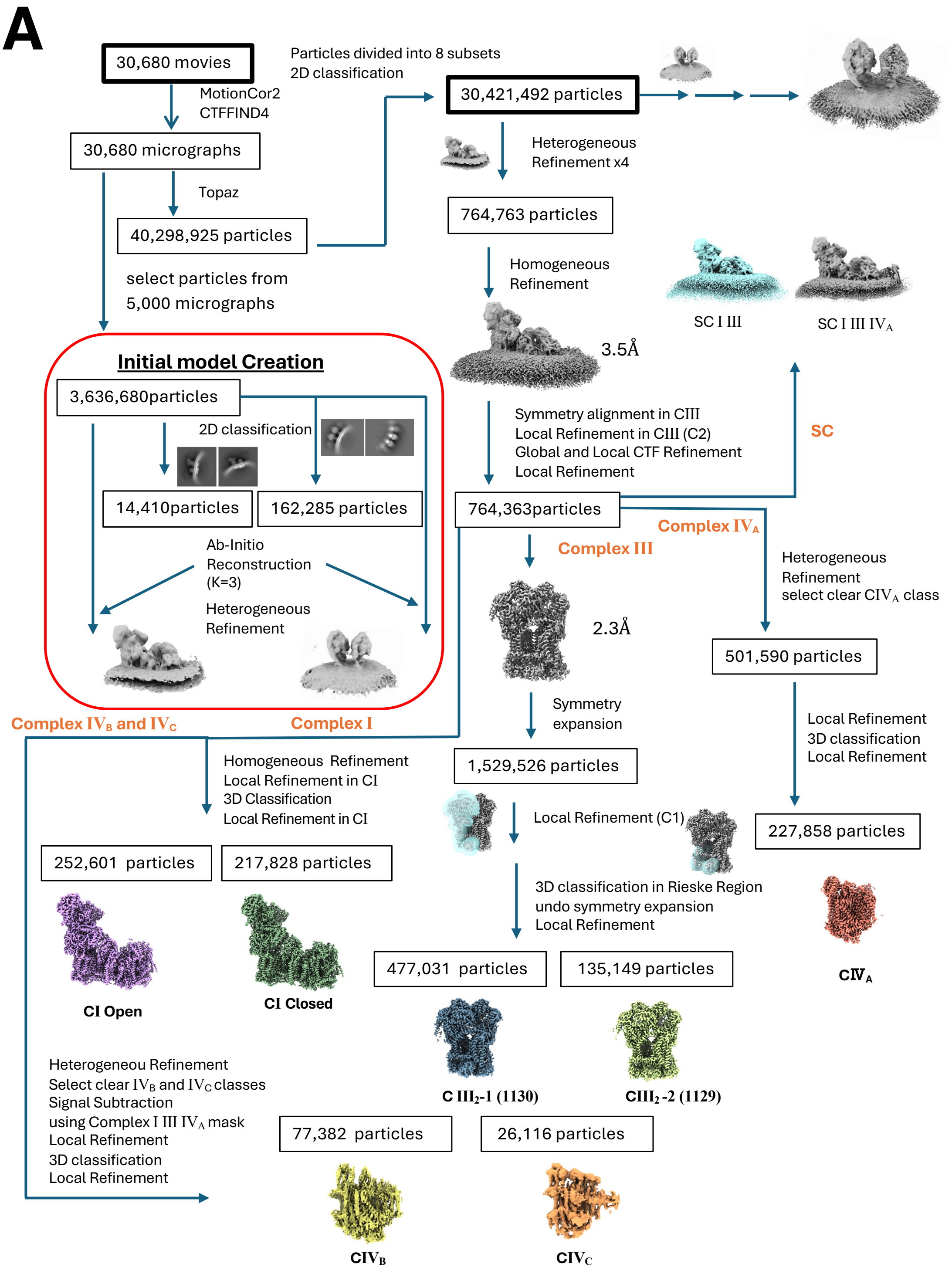

A
30,680 movies
MotionCor2
CTFFIND4
30,680 micrographs
Topaz
40,298,925 particles
select particles from
5,000 micrographs
Particles divided into 8 subsets
2D classification
30,421,492 particles
Heterogeneous
Refinement x4
764,763 particles
Homogeneous
Refinement
3.5Å
SC I III
SC I III IVA
Initial model Creation
3,636,680particles
2D classification
14,410particles
162,285 particles
Ab-Initio
Reconstruction
(K=3)
Heterogeneous
Refinement
Symmetry alignment in CIII
Local Refinement in CIII (C2)
Global and Local CTF Refinement
Local Refinement
SC
764,363particles
Complex IVA
Complex III
Heterogeneous
Refinement
select clear CIVA class
2.3Å
501,590 particles
Complex IVB and IVC
Complex I
Symmetry
expansion
1,529,526 particles
Local Refinement
3D classification
Local Refinement
Homogeneous Refinement
Local Refinement in CI
3D Classification
Local Refinement in CI
Local Refinement (C1)
227,858 particles
252,601 particles
217,828 particles
3D classification in Rieske Region
undo symmetry expansion
Local Refinement
477,031 particles
135,149 particles
CIVA
CI Open
CI Closed
C III2-1 (1130)
CIII2 -2 (1129)
Heterogeneou Refinement
Select clear IVB and IVC classes
Signal Subtraction
using Complex I III IVA mask
Local Refinement
3D classification
Local Refinement
77,382 particles
26,116 particles
CIVB
CIVC

**B**

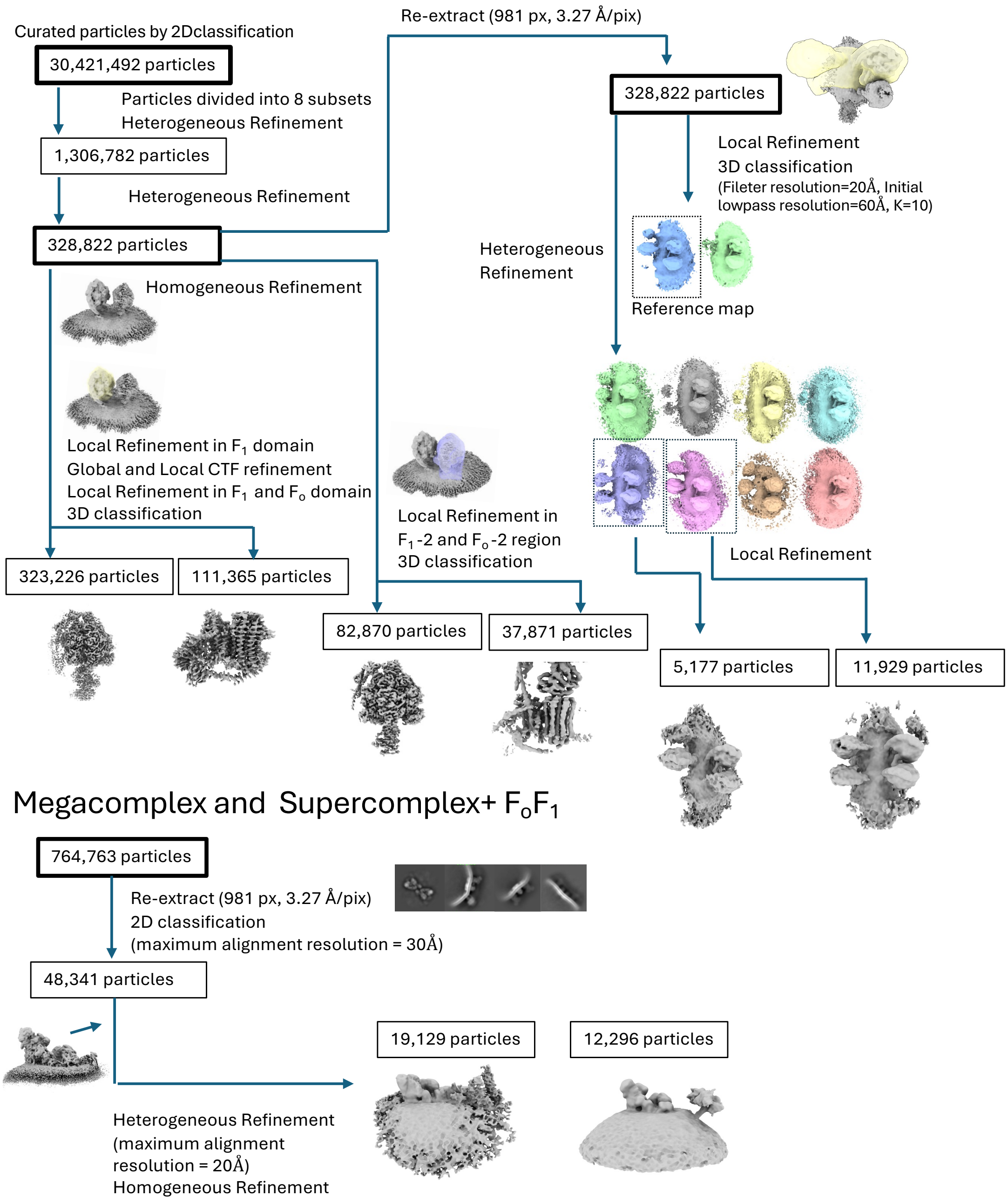
Curated particles by 2Dclassification
30,421,492 particles
Particles divided into 8 subsets
Heterogeneous Refinement
1,306,782 particles
Heterogeneous Refinement
328,822 particles
Re-extract (981 px, 3.27 Å/pix)
328,822 particles
Local Refinement
3D classification
(Fileter resolution=20Å, Initial lowpass resolution=60Å, K=10)
Heterogeneous Refinement
Reference map
Homogeneous Refinement
Local Refinement in F1 domain
Global and Local CTF refinement
Local Refinement in F1 and Fo domain
3D classification
Local Refinement in
F1-2 and Fo-2 region
3D classification
Local Refinement
323,226 particles
111,365 particles
82,870 particles
37,871 particles
5,177 particles
11,929 particles
Megacomplex and Supercomplex+ FoF1
764,763 particles
Re-extract (981 px, 3.27 Å/pix)
2D classification
(maximum alignment resolution = 30Å)
48,341 particles
19,129 particles
12,296 particles
Heterogeneous Refinement
(maximum alignment
resolution = 20Å)
Homogeneous Refinement

**C**

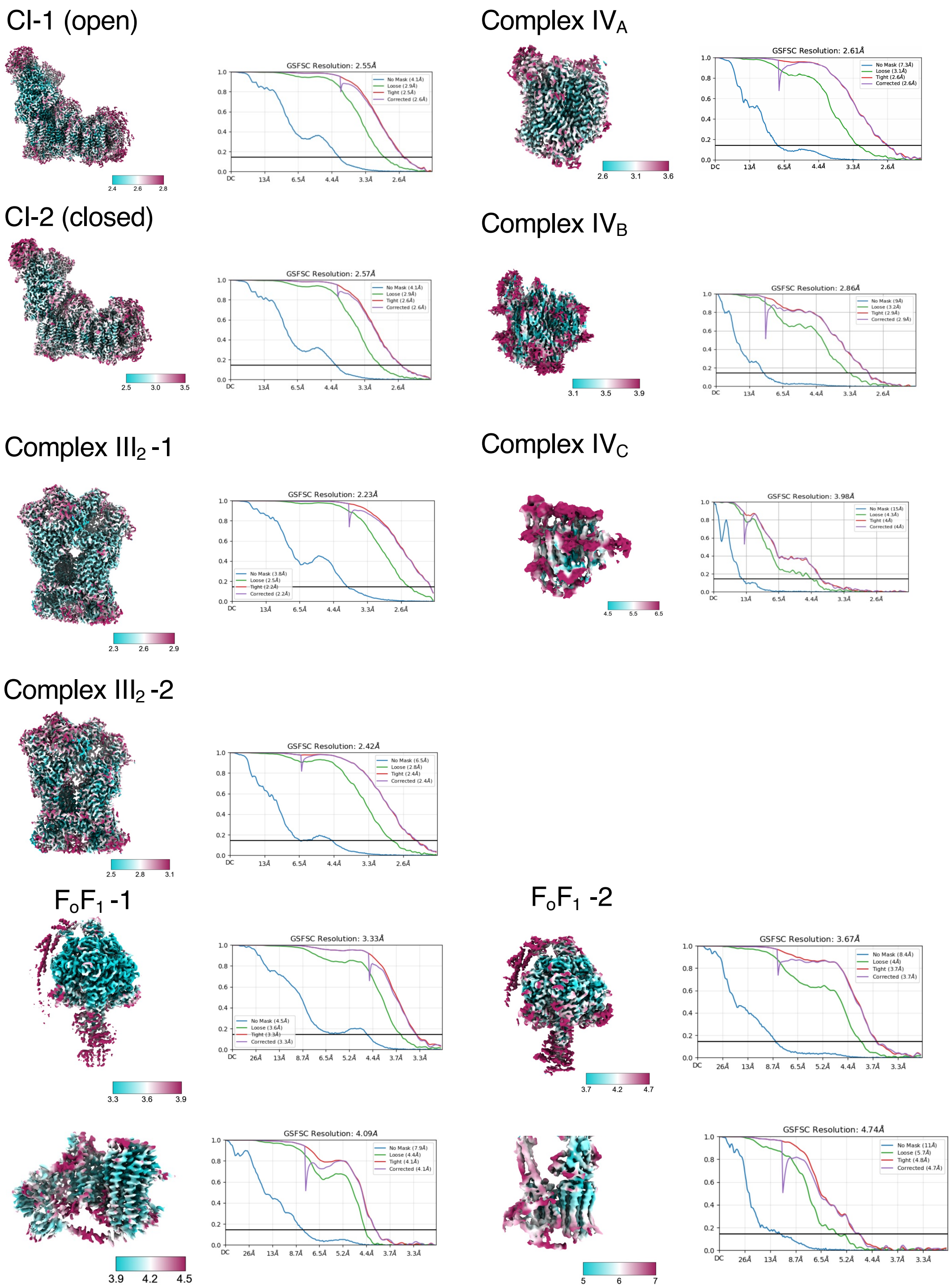

CI-1 (open)
GSFSC Resolution: 2.55Å
CI-2 (closed)
GSFSC Resolution: 2.57Å
Complex III2 -1
GSFSC Resolution: 2.23Å
Complex III2 -2
GSFSC Resolution: 2.42Å
Complex IVA
GSFSC Resolution: 2.61Å
Complex IVB
GSFSC Resolution: 2.86Å
Complex IVC
GSFSC Resolution: 3.98Å
FoF1 -1
GSFSC Resolution: 3.33Å
GSFSC Resolution: 4.09Å
FoF1 -2
GSFSC Resolution: 3.67Å
GSFSC Resolution: 4.74Å

**Figure S1. Analytical flowcharts and structural resolution of respiratory complexes.** (A) Workflow for the structural analysis of respiratory supercomplexes. (B) Workflow for the structural analysis of $F_oF_1$ ATP synthase oligomers. Detailed procedures are described in the *Materials and Methods* section. (C) Fourier Shell Correlation (FSC) curves for each complex. Local resolution maps are color-coded from blue (high resolution) to red (low resolution).

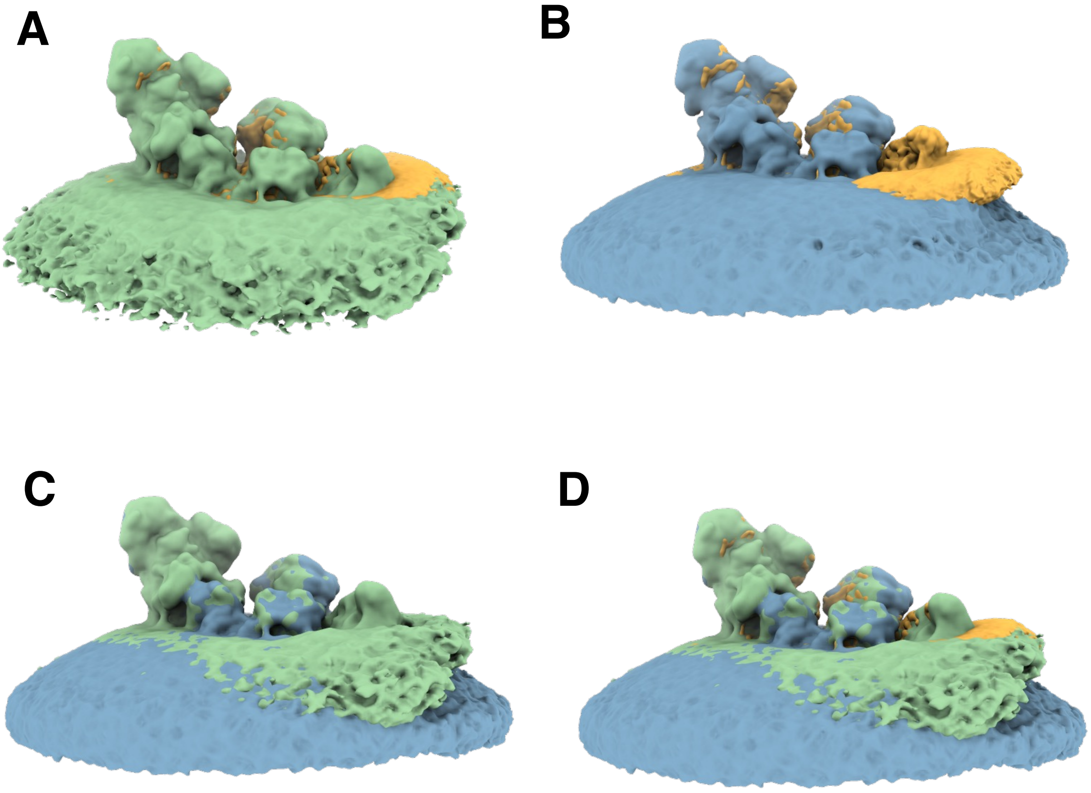


**Figure S2. SMPs membrane curvature influenced by complex IV composition.**
(A) Overlay of the cryo-EM maps of porcine mitochondrial $CI_1CIII_2CIV_1$ (yellow) and the present $CI_1CIII_2CIV_3$ complex. (B) Overlay of porcine mitochondrial $CI_1CIII_2CIV_1$ (yellow) and the present $CI_1CIII_2$ complex. (C) Overlay of the present $CI_1CIII_2$ and $CI_1CIII_2CIV_3$ cryo-EM maps. (D) Combined overlay of all three cryo-EM maps.

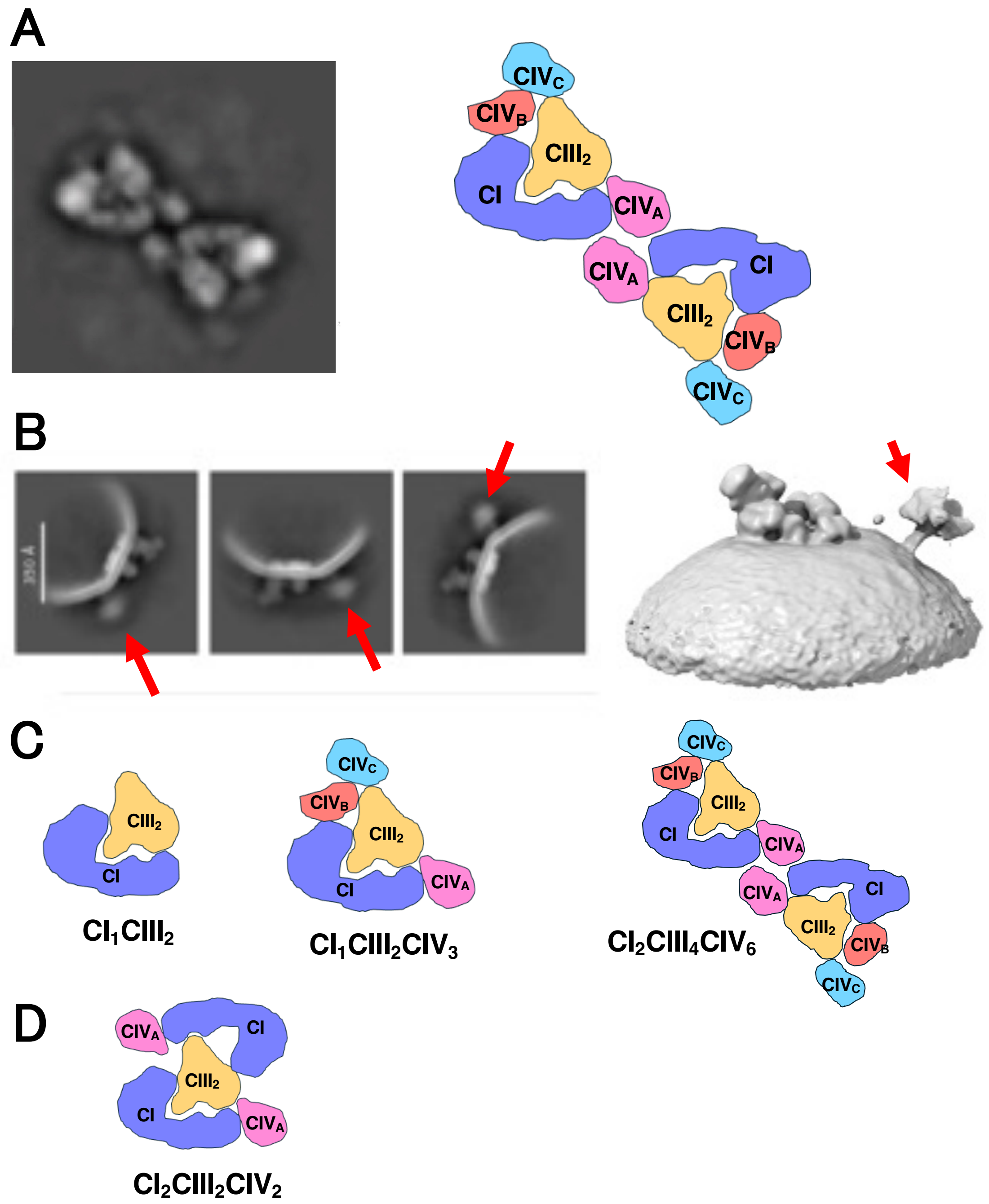


**Figure S3. Mega complexes observed on SMPs.**

(A) Representative 2D class averages (left) and schematic diagram (right) of the newly identified $CI_2CIII_4CIV_6$ mega complex. (B) 2D class averages (left) and 3D reconstruction (right) of supercomplexes (SCs) containing visible $F_1$ heads indicated by red arrows. (C) Schematic representation of the complex composition of the newly identified SCs. (D) $CI_2CIII_2CIV_2$ Mega complex previously identified in porcine mitochondria.

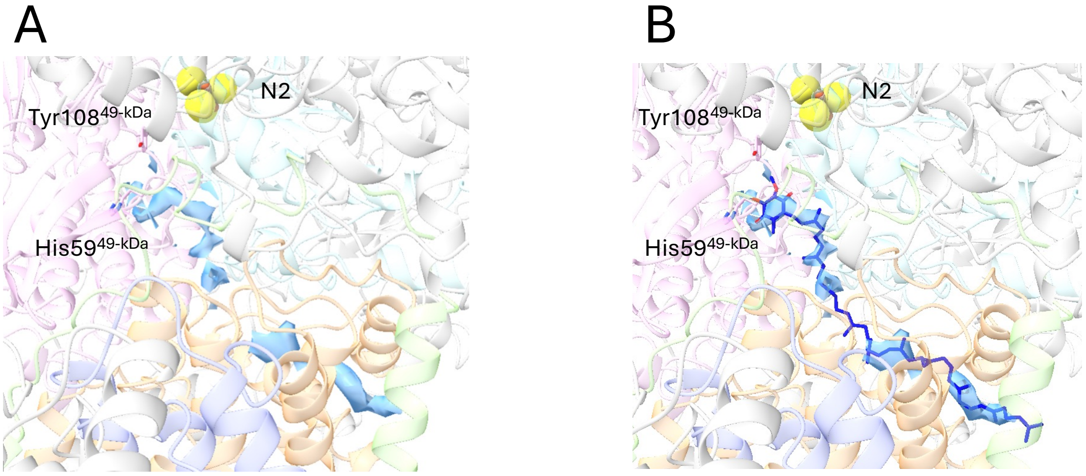


**Figure S4. Quinone tunnel in complex I.** (A) Cryo-EM density corresponding to the quinone-binding tunnel is highlighted in blue. (B) A model of ubiquinone-10 (Q-10) is fitted into the tunnel region.

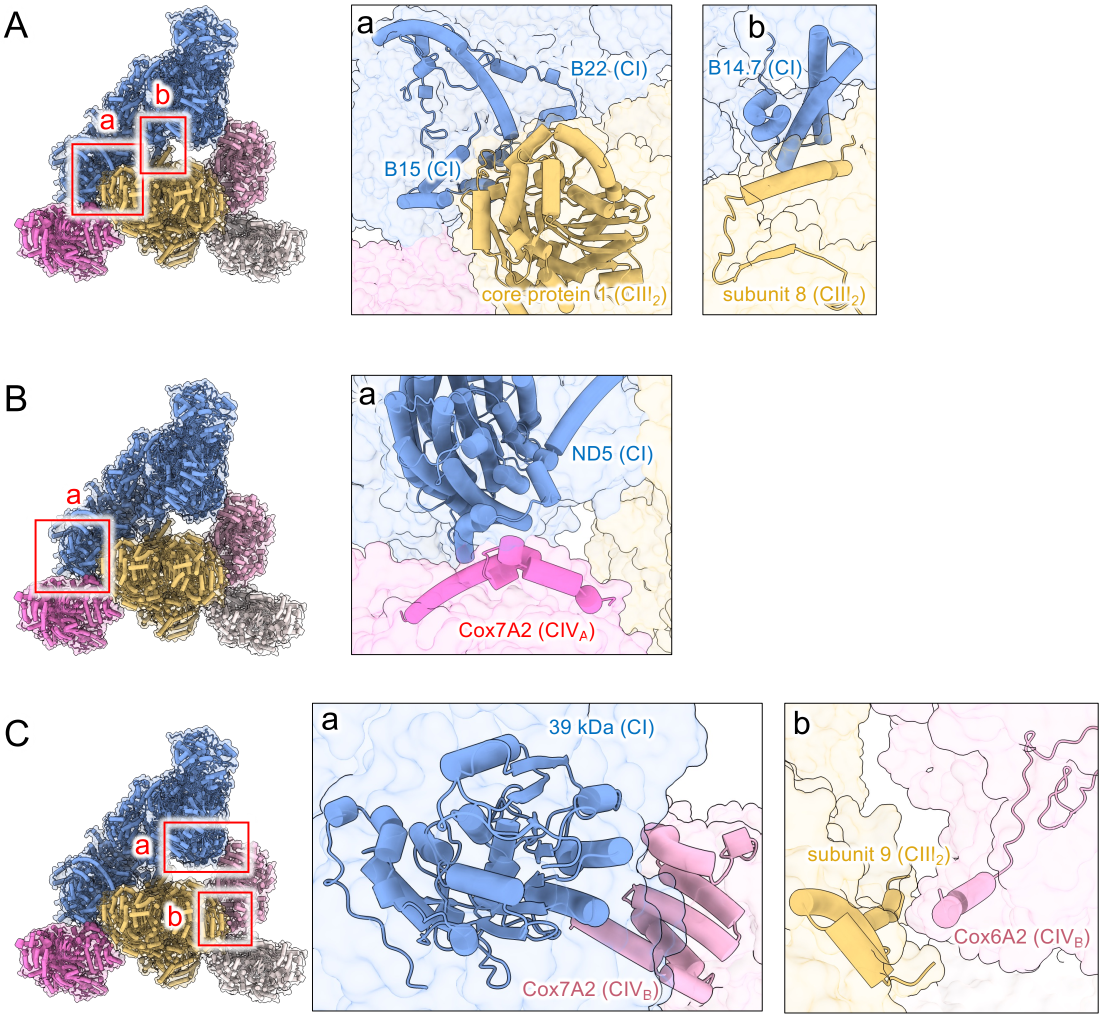


**Figure S5. Summary of the contact sites between respiratory complexes in the supercomplex (SC).**

(A) Contact sites between CI and $CIII_2$. $CIII_2$ associates with CI via (a) interactions between CI-B15 (NDUFB4) and CI-B22 (NDUFB9) with $CIII_2$–core protein 1 on the matrix side, and (b) interactions between CI-B14.7 (NDUFA11) and $CIII_2$–subunit 8 within the membrane bilayer. (B) Contact site between CI and $CIV_A$. $CIV_A$ interacts with CI through (a) contacts between the CI–ND5 subunit and the $CIV_A$–Cox7A subunit within the membrane bilayer. (C) Contact sites between $CIV_B$, CI, and $CIII_2$. $CIV_B$ is located in the interspace between CI and $CIII_2$, forming (a) a contact between CI–39-kDa and $CIV_B$–Cox5A on the matrix side, and (b) a contact between $CIII_2$–subunit 9 and $CIV_B$–Cox6A2 within the membrane bilayer.

### Table S1. Statistical Data on the Structures of Respiratory Complexes within SCs

| class | complex I | | complex III | | complex IV | | |
|---|---|---|---|---|---|---|---|
| | **state 1 (C1-open)** | **state 2 (C1-closed)** | **state 1 ($CIII_2$-1)** | **state 2 ($CIII_2$-2)** | **site A ($CIV_A$)** | **site B ($CIV_B$)** | **site C ($CIV_C$)** |
| EMDB ID | 65580 | 65581 | 65583 | 65584 | 65585 | 65586 | 65587 |
| PDB ID | 9W2U | 9W2V | 9W2X | 9W2Y | 9W2Z | - | - |
| **Data collection and processing** | | | | | | | |
| Magnification | 88,000 | 88,000 | 88,000 | 88,000 | 88,000 | 88,000 | 88,000 |
| Voltage(kV) | 300 | 300 | 300 | 300 | 300 | 300 | 300 |
| Microscope | Titan Krios | Titan Krios | Titan Krios | Titan Krios | Titan Krios | Titan Krios | Titan Krios |
| Total dose ($e^-$/Å $^2$) | 50 | 50 | 50 | 50 | 50 | 50 | 50 |
| Pixel size( Å /pix) | 0.84 | 0.84 | 0.84 | 0.84 | 0.84 | 0.84 | 0.84 |
| Defocus range(μm) | -0.8 to -2.0 | -0.8 to -2.0 | -0.8 to -2.0 | -0.8 to -2.0 | -0.8 to -2.0 | -0.8 to -2.0 | -0.8 to -2.0 |
| symmetry imposed | C1 | C1 | C1 | C1 | C1 | C1 | C1 |
| Initial particle | 40,298,925 | 40,298,925 | 40,298,925 | 40,298,925 | 40,298,925 | 40,298,925 | 40,298,925 |
| Final Particle | 252,601 | 217,828 | 477,031 | 135,149 | 227,858 | 77,382 | 26,116 |
| Map resolution( Å ) | 2.6 | 2.6 | 2.2 | 2.4 | 2.61 | 2.86 | 3.98 |
| FSC threshold | 0.143 | 0.143 | 0.143 | 0.143 | 0.143 | 0.143 | 0.143 |
| **Refinement** | | | | | | | |
| Initial model used | This study | This study | This study | This study | This study | This study | This study |
| Model resolution | 2.9 | 2.8 | 2.3 | | | | |
| FSC threshold | 0.5 | 0.5 | 0.5 | 0.5 | 0.5 | 0.5 | 0.5 |
| **Model composition** | | | | | | | |
| Nonhydrogen atoms | 68,035 | 68,568 | 33,266 | 33,266 | 14,833 | - | - |
| Protein residues | 8221 | 8289 | 4206 | 4206 | 1818 | - | - |
| Ligands | 1K, 1FMN, 1MG, 1CHD, 6CDL, 6SF4, 1PC1, 1GTP, 1ZN, 2FES, 1NDP, 1MYR, 18-3PE | 1K, 1FMN, 1MG, 6CDL, 6SF4, 1PC1, 1GTP, 1ZN, 2FES, 1NDP, 19-3PE | 6HEM, 2FES | 6HEM, 2FES | 1NA, 1MG, 1CU, 1CUA, 1PER, 1PGV, 1ZN, 2HEA, 1PEK | - | - |
| **R.m.s deviations** | | | | | | | |
| Bond length (Å) | 0.004 | 0.007 | 0.006 | 0.003 | 0.006 | - | - |
| Bond Angles (°) | 0.788 | 0.846 | 0.991 | 1.709 | 1.206 | - | - |
| **Validation** | | | | | | - | - |
| MolProbity score | 1.61 | 1.74 | 1.77 | 1.75 | 2.25 | - | - |
| Clashscore | 4.75 | 6.09 | 3.85 | 3.46 | 8.71 | - | - |
| Rotamer outlier (%) | 2.16 | 2.31 | 4.14 | 3.06 | 3.86 | - | - |
| CaBALM outlier (%) | 1.47 | 1.31 | 1.92 | 2.33 | 2.67 | - | - |
| **Ramachandran plot** | | | | | | | |
| Favored (%) | 97.46 | 97.32 | 97.31 | 96.35 | 95.25 | - | - |
| Allowed (%) | 2.43 | 2.64 | 2.67 | 3.65 | 4.75 | - | - |
| Disallowed (%) | 0.1 | 0.04 | 0.02 | 0.00 | 0.00 | - | - |

**Table S2. Statistical Data on the Structures of $F_1$ and $F_o$ domain within $F_oF_1$ oligomers**

| | $F_oF_1$-1 | | $F_oF_1$-2 |
|---|---|---|---|
| | F1 | Fo | F1 |
| EMDB ID | 65577 | 65578 | 65579 |
| PDB ID | 9W2R | 9W2S | 9W2T |
| **Data collection** | | | |
| Magnification | 88,000 | 88,000 | 88,000 |
| Voltage(kV) | 300 | 300 | 300 |
| Microscope | Titan Krios | Titan Krios | Titan Krios |
| Total dose ($e^-$/ Å $^2$) | 50 | 50 | 50 |
| Pixel size( Å /pix) | 0.84 | 0.84 | 0.84 |
| Defocus range(μm) | -0.8 to -2.0 | -0.8 to -2.0 | -0.8 to -2.0 |
| symmetry imposed | C1 | C1 | C1 |
| Initial particle | 30,421,492 | 30,421,492 | 30,421,492 |
| Final Particle | 328,822 | 111,365 | 328,822 |
| Map resolution( Å ) | 3.4 | 4.1 | 4.0 |
| FSC threshold | 0.143 | 0.143 | 0.143 |
| **Refinement** | | | |
| Initial model used | This study | This study | This study |
| Model resolution | 3.8 | 4.7 | 4.4 |
| FSC threshold | 0.5 | 0.5 | 0.5 |
| **Model composition** | | | |
| Nonhydrogen atoms | 39,135 | 9990 | 39,135 |
| Protein residues | 5135 | 1311 | 5155 |
| Ligands | 5MG, 4ATP, 1ADP | - | - |
| **R.m.s deviations** | | | |
| Bond length (Å) | 0.004 | 0.007 | 0.004 |
| Bond Angles (°) | 0.713 | 1.148 | 0.871 |
| **Validation** | | | |
| MolProbity score | 1.58 | 1.51 | 1.70 |
| Clashscore | 8.27 | 5.94 | 10.09 |
| Rotamer outlier (%) | 0.12 | 0.20 | 0.12 |
| CaBALM outlier (%) | 1.71 | 1.22 | 1.47 |
| **Ramachandran plot** | | | |
| Favored (%) | 97.29 | 96.89 | 96.98 |
| Allowed (%) | 2.71 | 3.11 | 3.02 |
| Disallowed (%) | 0.00 | 0.00 | 0.00 |
